# FMRFa receptor stimulated Ca^2+^ signals alter the activity of flight modulating central dopaminergic neurons in *Drosophila melanogaster*

**DOI:** 10.1101/336768

**Authors:** Preethi Ravi, Deepti Trivedi, Gaiti Hasan

## Abstract

Neuropeptide signaling influences animal behavior by modulating neuronal activity and thus altering circuit dynamics. Insect flight is a key innate behavior that very likely requires robust neuromodulation. Cellular and molecular components that help modulate flight behavior are therefore of interest and require investigation. In a genetic RNAi screen for G-protein coupled receptors that regulate flight bout durations, we earlier identified several receptors, including the receptor for the neuropeptide FMRFa (FMRFaR). To further investigate modulation of insect flight by FMRFa we generated CRISPR-Cas9 mutants in the gene encoding the *Drosophila* FMRFaR. The mutants exhibit significant flight deficits with a focus in dopaminergic cells. Expression of a receptor specific RNAi in adult central dopaminergic neurons resulted in progressive loss of sustained flight. Further, genetic and cellular assays demonstrated that FMRFaR stimulates intracellular calcium signaling through the IP_3_R and helps maintain neuronal excitability in a subset of dopaminergic neurons for positive modulation of flight bout durations.

**Author summary:** Neuropeptides play an important role in modulating neuronal properties such as excitability and synaptic strength and thereby influence innate behavioral outputs. In flying insects, neuromodulation of flight has been primarily attributed to monoamines. In this study, we have used the genetically amenable fruit fly, *Drosophila melanogaster* to identify a neuropeptide receptor that is required in adults to modulate flight behavior. We show from both knockdown and knockout studies that the neuropeptide receptor, *FMRFaR*, present on a few central dopaminergic neurons, modulates the duration of flight bouts. Overexpression of putative downstream molecules, the IP_3_R, an intracellular Ca^2+^-release channel, and CaMKII, a protein kinase, significantly rescue the flight deficits induced by knockdown of the *FMRFaR*. Our data support the idea that FMRFaR and CaMKII help maintain optimal membrane excitability of adult dopaminergic neurons required to sustain longer durations of flight bouts. We speculate that the ability to maintain longer flight bouts in natural conditions enhances the individual’s capacity to search and reach food sources as well as find sites suitable for egg laying.

## Introduction

Neuromodulation of animal behavior by neuropeptides is ubiquitous among vertebrates and invertebrates (1, 2). Unlike fast acting neurotransmitters, neuropeptides and their receptors influence neuronal activity and circuit dynamics by modulating presynaptic neurotransmitter release. The mechanisms for doing this include changes in ion channel and transporter function as well as regulation of gene expression (1, 2). The neural action of neuropeptides can either be local or at long distances by release into circulation and can influence intrinsic behaviors such as feeding, mating, sleep and aggression (3, 4). An important and critical behavior in flying insects is flight. Altered flight behavior, in particular the inability to maintain long durations of flight bouts, can impinge on the fly’s ability to optimally search and reach food sources as well as sites suitable for egg laying. Neuromodulation of insect flight has thus far been attributed primarily to biogenic amines (5–7). A role for neuropeptide-based modulation of flight behavior has remained largely unexplored.

In invertebrates, neuropeptides activate G-protein coupled receptors (GPCRs) (8) followed by generation of soluble second messengers such as cAMP (9–12) or inositol 1,4,5-trisphosphate (IP_3_) (13–15). IP_3_ binds to and activates the endoplasmic reticulum (ER) localized Ca^2+^ channel, the IP_3_ receptor (IP_3_R), resulting in release of calcium from ER-stores (16). Invertebrate neuropeptide receptors that stimulate IP_3_-mediated Ca^2+^ release include the Pigment Dispersing Factor Receptor (PdfR) (14), FMRFaR (13–15) amongst others (8, 17). We identified the PdfR and FMRFaR in a genetic screen for neuronal GPCRs that regulate flight in *Drosophila* through IP_3_-mediated Ca^2+^ release (14). Further investigation of PdfR demonstrated a role for this receptor in both the developing and adult flight circuit, though the identity of Pdf responsive neurons that affect flight remained ambiguous (14). FMRFaR has been described earlier in the context of an escape response to intense light in *Drosophila* larvae (13) and the larval to pupal transition under nutrient-limiting conditions (15). Adult behaviors implicating the FMRFaR include startle-induced locomotor activity (18) and adaptive sleep following heat stress (19).

In the context of flight behavior, a neuronal requirement for IP_3_-mediated Ca^2+^ release was initially described in flight deficient IP_3_R mutants (20). Subsequent cellular and molecular studies identified a role for IP_3_-mediated Ca^2+^ release in dopaminergic neurons during pupal stages, when the flight circuit matures (7). Interestingly though, flight deficits were also observed upon temporal knockdown of the *FMRFaR* exclusively in mature neurons (14). Thus far, flight deficits arising from adult specific reduction of IP_3_/Ca^2+^ signaling have remained unexplored. Here we have investigated if FMRFaR mediated Ca^2+^ signaling is required for *Drosophila* flight by generating new CRISPR-Cas9 mediated mutants for the *FMRFaR*. Flight deficits arising from specific knockdown in adult dopaminergic neurons suggests a role for FMRFaR in modulating adult flight and implicate the IP_3_R and possibly CaMKII as downstream signaling components. Genetic and cellular assays indicate that the FMRFaR on adult dopaminergic neurons helps maintain optimal membrane excitability which could potentially be required for synaptic release of dopamine.

## Results

### FMRFaR is required in dopaminergic neurons for maintenance of flight

The neuropeptide receptor, FMRFaR, was identified amongst other G-protein coupled receptors (GPCRS) as a positive regulator of *Drosophila* flight in a pan-neuronal screen, where genetic data supported IP_3_R mediated Ca^2+^ signaling and store-operated calcium entry (SOCE) as the down-stream effectors of receptor activation (14). We confirmed these initial observations by RNAi mediated knockdown of the *FMRFaR* with another pan-neuronal GAL4 (*nSybGAL4*) in the central nervous system (CNS), followed by measurement of flight bout durations in tethered flies in response to a gentle air-puff. Unlike previous measurements where flight times were recorded for a maximum of 30 seconds (14), in the current study we monitored flight for 15 minutes (900 seconds), which allows for resolution of longer flight bout durations. Upon *FMRFaR* knockdown in *nSyb* positive neurons, significantly shorter flight bouts were observed as compared to controls (*FMRFaR^RNAi^/+, nSyb/+*; Fig 1A). The efficacy of *FMRFaR* knockdown by the RNAi strain was confirmed in the adult CNS by pan-neuronal expression of the RNAi with *nSybGAL4*. A significant reduction in *FMRFaR* mRNA levels was observed (Fig 1B).

**Fig 1.**
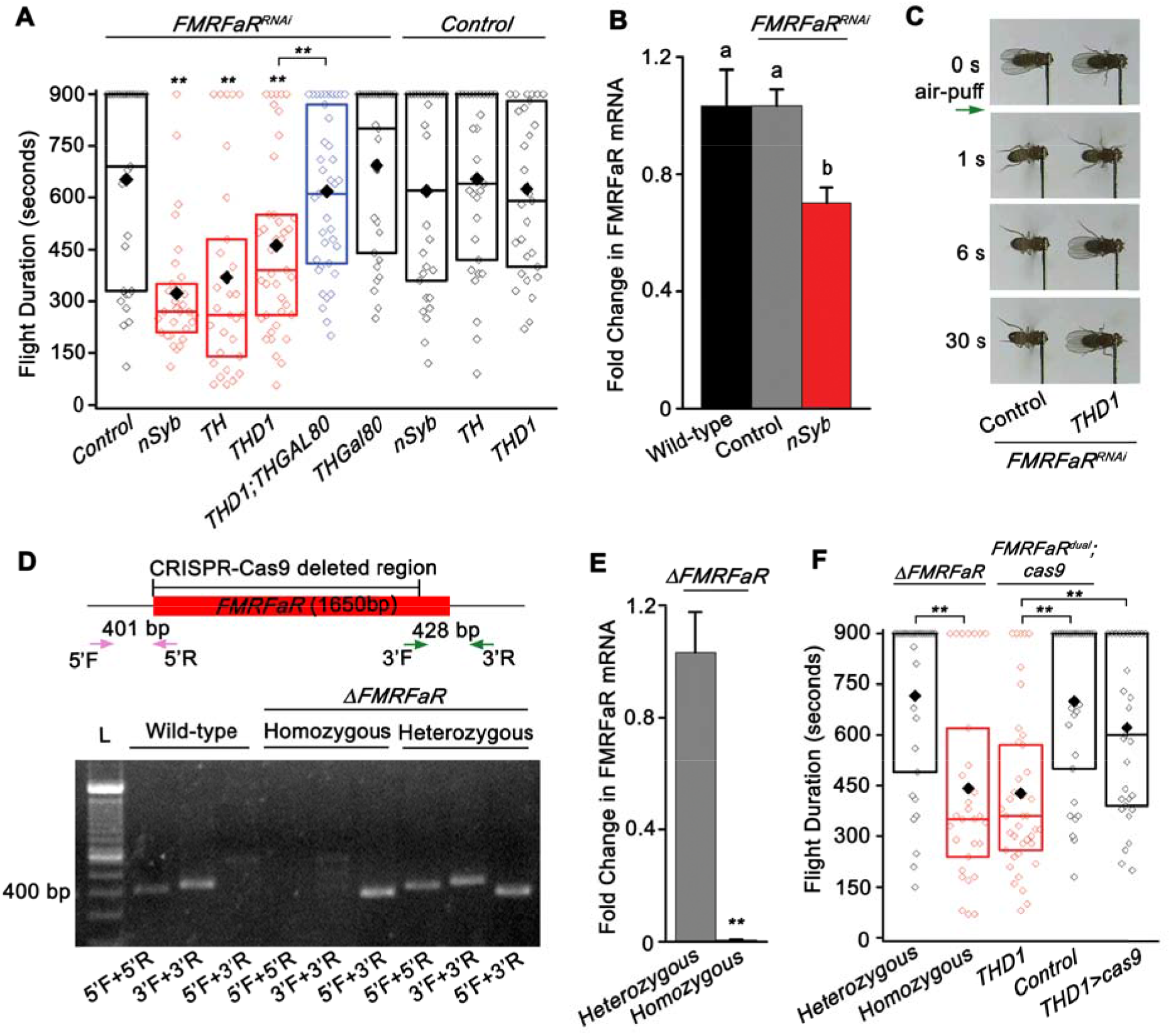
FMRFaR on dopaminergic neurons is required for flight. (A) Box plot showing duration of flight bouts in flies expressing *FMRFaR^RNAi^* in the indicated neuronal domains (red). In every box plot, the limits extend from 25^th^ to 75^th^ percentile, the line and solid diamond represent the median and mean respectively and the open diamonds show the individual data points. Significance was calculated by comparing with either the control genotypes (*FMRFaR^RNAi^/+, nSyb/+, TH/+, THD1/+*) or as indicated by horizontal lines above the graph (n≥30, **p<0.01, Mann-Whitney U-test). Genotypic controls have been obtained throughout by crossing the strains to be tested with wild-type, *Canton-S*. The ‘+’ in all control genotypes denotes the wild-type allele. (B) Quantitative PCR validation of *FMRFaR* knockdown in *nSyb* positive neurons. Each bar represents mean of normalized fold change ± SEM (n—6, One-way ANOVA followed by post-hoc Tukey’s test; the same alphabet above each bar represents statistically indistinguishable groups; different alphabet represents p<0.01). (C) Snapshots of flight videos of the indicated genotypes (*FMRFaR^RNAi^/+ and THD1>FMRFaR^RNAi^*) at time points before and after air-puff stimulus (green arrow). Flies of the control genotype continued to fly for time periods beyond those shown in the images. (D) Representation of the *FMRFaR* gene locus. The CRISPR-Cas9 based deletion removes ~1500 bp of the 1650 bp coding sequence. Primers used to amplify the 5’ (pink: 5’F, 5’R) and 3’ (green: 3’F, 3’R) ends of the gene are indicated by arrows. Agarose gel showing PCR products from wild-type, heterozygous and homozygous knockouts of *FMRFaR*. L denotes DNA Ladder. (E) Quantitative PCR validation of *FMRFaR* CRISPR knockout from adult brains (n≥3, **p<0.01, unpaired t-test). (F) Flight bout duration of flies with CRISPR-Cas9 based knockout of *FMRFaR*. Statistical comparisons are as indicated by horizontal lines (n≥30, **p<0.01, Mann-Whitney U-test).

Similar to the pan-neuronal knockdown of *FMRFaR*, flight deficits were also observed in flies with *FMRFaR^RNAi^* targeted to a large number of dopamine-synthesizing cells driven by *THGAL4* (Fig 1A). Because non-neuronal expression of *THGAL4* has been documented (21), we tested GAL4s with restricted expression in neurons (22). For this, we used the *THD* and *THCGAL4* driver lines that mark non-overlapping subsets of central dopaminergic neurons (22). Knockdown of *FMRFaR* using either the *THD1GAL4* or the *THD’GAL4* resulted in flight deficits that were equivalent to the phenotype observed with *THGAL4-*driven knockdown (Fig 1A, C, S1A Fig; *TH>FMRFaR^RNAi^* vs. *THD1>FMRFaR^RNAi^* or *TH>FMRFaR^RNAi^* vs *THD’>FMRFaR^RNAi^* are not significantly different from each other; p>0.05; Mann-Whitney U-test). However, the *THDGAL4* strains are also known to express in some non-dopaminergic neurons (22). To test the role, if any, of FMRFaR function in non-dopaminergic neurons marked by *THDGAL4s*, expression of the *FMRFaR^RNAi^* was blocked specifically in TH-expressing neurons with the *THGAL80* transgene (23). Importantly, the flight deficits observed in *THD>FMRFaR^RNAi^* flies were reversed upon introduction of *THGAL80* confirming that such flight deficits arose exclusively from dopaminergic neurons marked by these GAL4s (Fig 1A; blue bar, S1A Fig; blue bar). On the other hand, flies with knockdown of the *FMRFaR* using the *THC1* or *THC’GAL4s* did not result in significant flight deficits, indicating that dopaminergic neurons unique to *THD1* or *THD’GAL4* require FMRFaR mediated signaling to regulate flight bout durations (S1A Fig). *THDGAL4* strains express in central dopaminergic neurons comprising the Protocerebral Posterior Lateral 1 (PPL1) and Protocerebral Posterior Medial 3 (PPM3) clusters and these two neuronal clusters are not marked by the *THC’GAL4* (7, 22). Furthermore, the reduced flight deficit observed with *FMRFaR* knockdown in dopaminergic neurons was not a consequence of overall reduction in motor activity because *THD1>FMRFaR^RNAi^* flies showed normal locomotor activity as compared to their genetic controls (S1B Fig). Having identified the dopaminergic subsets where FMRFaR is required for flight, subsequent experiments were performed with *THD1GAL4* as the dopaminergic expression driver.

RNAi knockdown of *FMRFaR* with *nSybGAL4* reduced *FMRFaR* levels by about 30% (Fig 1B). To further refine our understanding of the FMRFaR’s role in *Drosophila* flight, a complete knockout of the *FMRFaR (ΔFMRFaR*) was generated by the CRISPR-Cas9 method (24). The strategy employed, removed nearly 1.5 kb of the 1.6 kb coding region of *FMRFaR* (Fig 1D). Primers spanning the *FMRFaR* locus (5’F+3’R) amplified only about ~400bp genomic fragment from *FMRFaR* homozygous and heterozygous knockouts, thereby confirming *FMRFaR* gene knockout (Fig 1D). Other primer pair combinations (5’F+5’R and 3’F+3’R) helped us distinguish between the homozygotes and heterozygotes. Homozygous knockouts of the *FMRFaR* showed near complete loss of *FMRFaR* mRNA as well as reduced flight bout durations as compared to the heterozygotes (Fig 1E, F).

Next, we tested dopaminergic neuron specific knockout of *FMRFaR* by expression of *UAScas9* in the genetic background of flies with ubiquitous expression of guide RNAs that target the gene for *FMRFaR (THD1>FMRFaR^dual^;cas9*; see Materials and methods). Flies with knockout of the *FMRFaR* in *THD1* cells flew for significantly shorter durations as compared to the appropriate genetic controls (*FMRFaR^dual^/+;cas/+* - Fig 1F; 4^th^ bar in black and *THD1/+* - Fig 1A; last bar in black; p<0.01; Mann-Whitney U-test) (Fig 1F). To account for non-specific effects of *cas9* expression, we tested flies expressing only the *cas9* transgene in *THD1* neurons (*THD1>cas9*). The flight deficit observed with *FMRFaR* knockout in *THD1* cells was significantly different from *THD1>cas9* flies (Fig 1F; *THD1>cas9* vs *THD1>FMRFaR^dual^;cas9*; p<0.01; Mann-Whitney U-test). The near identical flight deficits observed between *FMRFaR* null flies and flies with *FMRFaR* knockout in *THD1* cells suggests that modulation of flight bout duration by FMRFaR may derive solely from dopaminergic neurons (Fig 1F; *THD1>FMRFaR^dual^;cas9* vs *ΔFMRFaR* Homozygous; p>0.05; Mann-Whitney U-test). To test this idea, we next investigated whether *FMRFaR* is enriched in dopaminergic neurons of the brain.

### FMRFaR is present on dopaminergic neurons and is functionally active

Owing to the lack of either an antibody for FMRFaR or GAL4 strains that reliably mark FMRFaR expressing neurons, we chose to employ molecular and functional assays to confirm the presence of FMRFaR on dopaminergic neurons. Towards this end, we sorted adult brain dopaminergic neurons marked by *THGAL4* driven *eGFP*, using fluorescence-activated cell sorting (FACS) and measured transcript levels of a few selected genes. Two markers of dopaminergic neurons, *ple* (encoding the enzyme Tyrosine Hydroxylase or TH) and *dDAT* (encoding a dopamine transporter) were highly expressed in GFP +ve cells, confirming homogeneity of the sorted population (Fig 2A). *FMRFaR* expression, as measured by qPCR, was significantly enriched in *TH* expressing neurons (GFP +ve), whereas expression of calcium signaling molecules like *dSTIM* and *dOrai* was similar in dopaminergic and non-dopaminergic neurons (Fig 2A).

**Fig 2.**
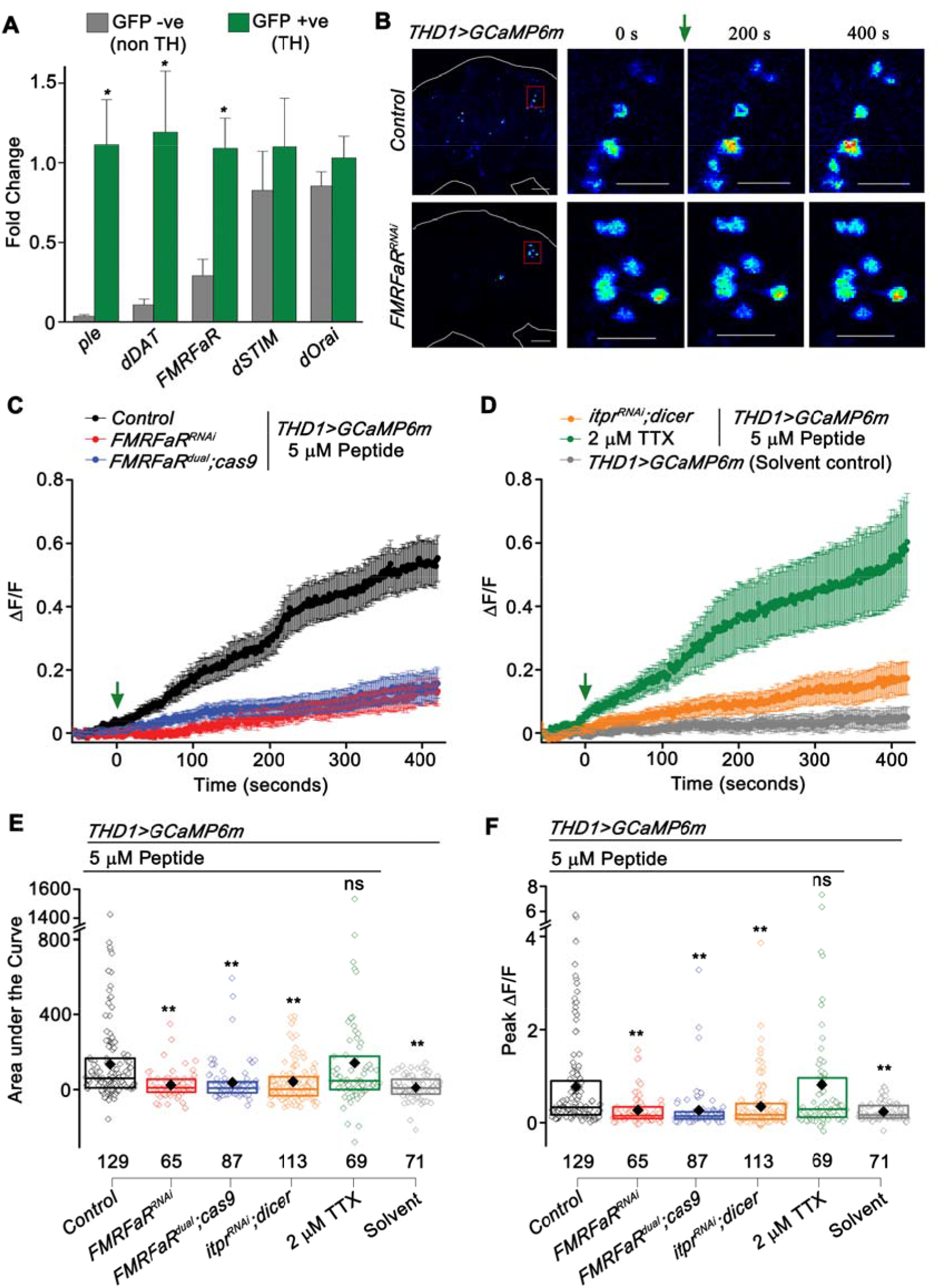
FMRFaR is expressed on dopaminergic neurons and is functionally active. (A) Quantitative PCR on FACS sorted dopaminergic (GFP +ve, TH) and non-dopaminergic (GFP-ve, non TH) neurons, shows enrichment of *FMRFaR* transcripts in dopaminergic neurons. Each bar represents normalized fold change as mean ± SEM (n≥4, *p<0.05, unpaired t-test) (B) Snap shots of GCaMP6m responses from the indicated genotypes. The red box represents the magnified region of the protocerebrum shown in subsequent images. Warmer colors denote increase in [Ca^2+^]. Scale bars indicate 20 μm. (C, D) Mean traces (±SEM) of normalized GCaMP6m fluorescence responses (ΔF/F) in *THD1GAL4* marked cells upon peptide or solvent addition (green arrow). (E) Area under the curve and (F) Peak ΔF/F quantified from 0 s to 420 s in (C, D). The box plot limits extend from 25^th^ to 75^th^ percentile. The line and solid diamond represent the median and mean respectively. Individual data points are shown as open diamonds. Numbers below indicate total number of cells imaged. Each bar is compared to the Control peptide stimulated condition shown in black (**p<0.01; ns - not significant; Mann-Whitney U-test).

The presence of active FMRFaR on central dopaminergic neurons was tested next. *THD1* neurons with expression of a genetically encoded calcium sensor, *GCaMP6m* (25) were tested for receptor activation in adult brain explants and calcium responses were specifically monitored from the PPL1 and PPM3 clusters, previously implicated in the regulation of flight bout durations (Fig 1A, S1A Fig) (7). Stimulation with one of the most abundantly expressed neuropeptide ligands of FMRFaR, DPKQDFMRFa (henceforth referred to as FMRFa) (26, 27), resulted in a slow calcium rise (Fig 2B, C, E, F). FMRFa peptide stimulated GCaMP6m response was significantly attenuated upon knockdown of the *FMRFaR* using either the RNAi or *FMRFaR^dual^;cas9* (Fig 2B, C, E, F). Peptide stimulation of *THD1* cells in the presence of a sodium channel inhibitor, 2 μM Tetrodotoxin (TTX) resulted in calcium responses that were comparable to that obtained in the absence of TTX (Fig 2D, E, F). This suggested that the rise in FMRFa-stimulated Ca^2+^ is due to direct activation of the FMRFaR and not a consequence of synaptic inputs from other neurons to the *THD1* neurons. Moreover, FMRFa stimulated Ca^2+^ signals attenuated significantly upon knockdown of *itpr* in *THD1* marked dopaminergic neurons (Fig. 2D, E, F). Taken together, these data suggest the presence of FMRFaR on central brain dopaminergic neurons and that receptor activation likely leads to Ca^2+^ release from the IP_3_R.

### FMRFaR is required in adult dopaminergic neurons for sustained flight

Pan-neuronal knockdown of the *FMRFaR* in an earlier study suggested a requirement of the receptor both during pupal development and in adult *Drosophila* neurons (14). To investigate the developmental stage(s) when FMRFaR signaling is required in dopaminergic neurons for maintaining flight, the TARGET (Temporal And Regional Gene Expression Targeting) system (28) was used for stage-specific *FMRFaR* knockdown. This system employs a temperature sensitive GAL80 element (*TubGAL80^ts^*) which represses GAL4 expression at 18°C. At 29°C the GAL80^ts^ is inactivated, thus allowing GAL4 driven expression of the UAS transgene. Flight durations of flies with *FMRFaR* knockdown in dopaminergic neurons throughout development (29°C) were reduced to less than 300 seconds as compared to genetic controls that on average flew for about 600 seconds (Fig 3A, S2A Fig). Flies with knockdown of the *FMRFaR* in larval stages displayed normal flight bouts (29°C Larval), whereas a modest reduction in flight bout durations was observed with pupal knockdown (29°C Pupal) (Fig 3A). Interestingly, knockdown of the *FMRFaR* in adult stages affected flight duration maximally, with exacerbation of the flight deficit over time, such that flies with knockdown for eight days flew for less than 150 seconds (29°C Adults) (Fig 3A). In contrast, the control genotypes maintained at 29°C for the same adult period could fly for an average of over 600 seconds (S2A Fig).

**Fig 3.**
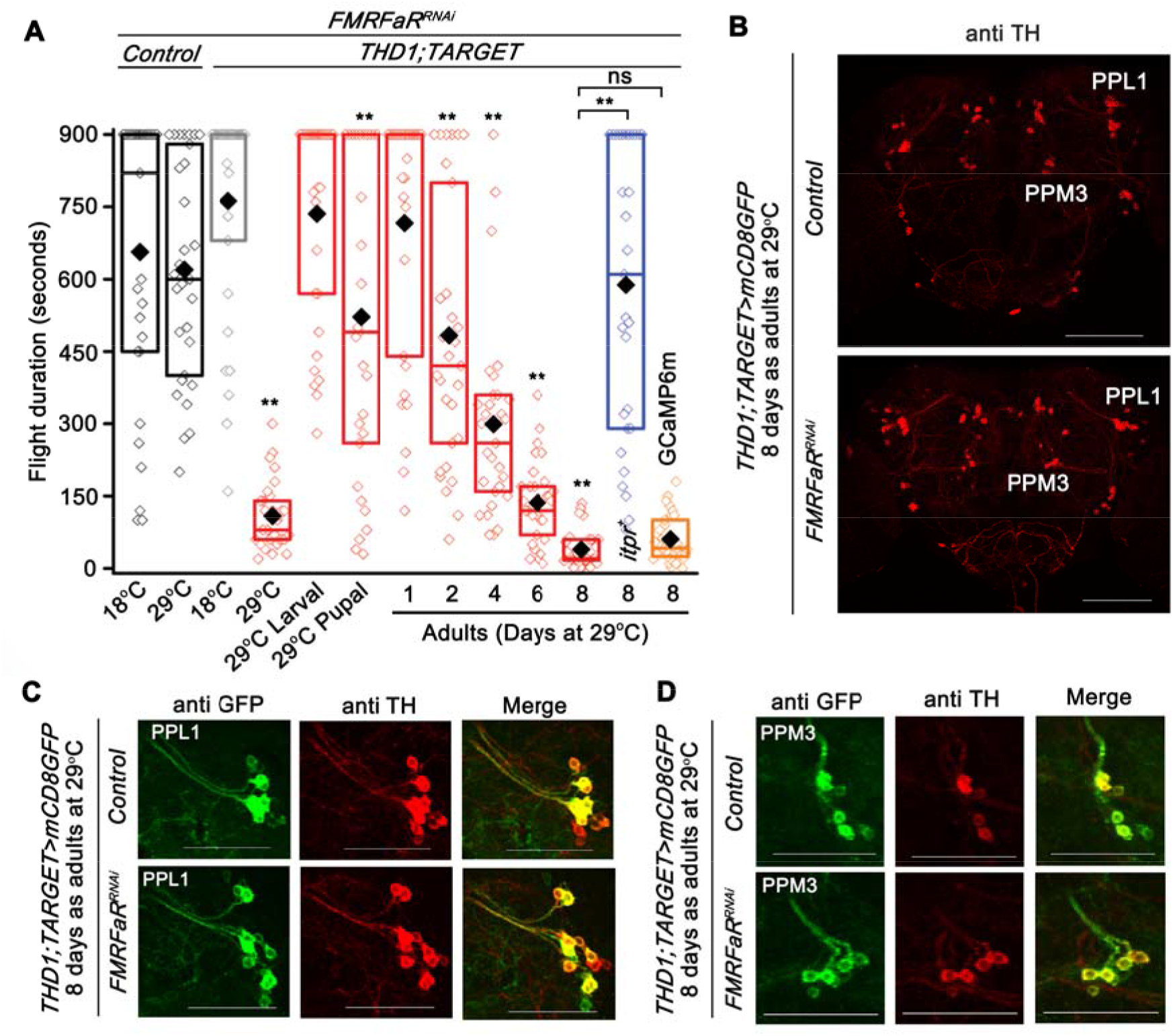
FMRFaR is required in adult dopaminergic neurons for sustained flight. (A) Box plot showing flight bout durations by temporal knockdown of *FMRFaR* in *THD1* neurons using the TARGET system (*THD1; TubGAL80^ts^>FMRFaR^RNAi^*), which allows effective expression of transgene at 29°C, but not at 18°C. Animals maintained at 18°C or 29°C throughout development and as adults are indicated as 18°C and 29°C, respectively. Knockdown of *FMRFaR* only during the larval, pupal or adult stages are shown as 29°C Larval, 29°C Pupal or 29°C Adult, respectively (red). Knockdown of *FMRFaR* in adult stages (29°C Adult) reduced flight bout durations significantly. The phenotype observed upon *FMRFaR* knockdown in adults for 8 days could be rescued by overexpression of the *itpr*^+^ transgene, but not with another UAS transgene, *GCaMP6m (THD1;TubGAL80^ts^>FMRFaR^RNAi^;itpr^+^* shown in blue and *THD1;TubGAL80^ts^>FMRFaR^RNAĩ^;GCaMP6m* shown in orange; n≥30, **p<0.01, ns - not significant; Mann-Whitney U-test). All comparisons are to the control condition, *THD1;TARGET>FMRFaR^RNAi^* at 18°C (grey bar; *THD1;TubGAL80^ts^>FMRFaR^RNAi^;* 18°C) or as indicated by horizontal lines above the graph (n≥30, **p<0.01, Mann-Whitney U-test). (B) Immunohistochemistry of representative adult brain samples corresponding to 8 day adult controls (above; *THD1;TubGAL80^ts^>mCD8GFP* Control) and *FMRFaR* knockdown (below; *THD1;TubGAL80^ts^>mCD8GFP;FMRFaR^RNAi^*). Anti-TH antibody was used to mark the various dopaminergic clusters, amongst which PPL1 and PPM3 clusters are indicated. Scale bars represent 100 μm. (C, D) Representative magnified views of the PPL1 (C) and PPM3 (D) clusters and their projections showing anti-GFP and anti-TH immunostaining in 8 day old adult brains of control (above; *THD1;TubGAL80^ts^>mCD8GFP* Control) and *FMRFaR* knockdown (below; *THD1;TubGAL80^ts^>mCD8GFP;FMRFaR^RNAi^*). Scale bar represents 50 μm.

Next, we tested the ability of IP_3_R overexpression to compensate for loss of *FMRFaR* in adult dopaminergic neurons. Indeed, flight deficits observed with *FMRFaR* knockdown in adults could be rescued to a significant extent by simultaneous overexpression of an *itpr*^+^ transgene (Fig 3A; blue bar; p<0.01; Mann-Whitney U-test), but not another *UAS* transgene, *GCaMP6m* (Fig 3A; orange bar; p>0.05; Mann-Whitney U-test). Rescue of *FMRFaR* knockdown by overexpression of *itpr*^+^ supports earlier findings that IP_3_R-mediated Ca^2+^ release functions downstream of FMRFaR (13) and is consistent with observations in Fig. 2D, E, F. Temporal and cell specific knockout of the *FMRFaR* with *FMRFaR^dual^* was attempted, but was not pursued because control flies with expression of *cas9* in adult *THD1;TARGET* cells exhibit flight deficits when maintained at 29°C for 8 days (S2A Fig, last lane in pink). Consistent adult specific phenotypes were observed with the *THD1GAL4* strain which has been used for subsequent experiments.

The cellular basis for flight deficits arising from loss of *FMRFaR* in *THD1* marked neurons was investigated next. Expression of endogenous TH and membrane-bound GFP driven by *THD1GAL4* was visualized in the PPL1 and PPM3 clusters of adult *Drosophila* brains at 8 days after knockdown of *FMRFaR*. Both TH and GFP immunoreactivity appeared as in a control genotype, and there was no apparent loss of dopaminergic neurons or their neurite projections (Fig 3B, C, D, S2B Fig). Taken together these data suggest that the FMRFaR is required in adult *THD1* neurons for sustaining flight bout durations. The increasing disability to maintain flight bouts for longer durations upon manipulation of *FMRFaR* levels in adult *THD1* neurons could likely be a consequence of increased RNAi expression over time.

### CaMKII in dopaminergic neurons is required for flight

Downstream signaling mechanisms that require FMRFaR activation in *THD1* dopaminergic neurons were investigated next. There is evidence demonstrating that FMRFa evoked modulation of excitatory junction potentials (EJPs) at *Drosophila* neuro-muscular junctions are dependent on the protein kinase, CaMKII (13, 29). Hence, we tested whether overexpression of wild-type *CaMKII (WT-CaMKII*) in dopaminergic cells rescued the deficit in flight bout durations observed upon reduced FMRFaR signaling. Overexpression of *WT-CaMKII* during pupal stages was insufficient to rescue flight deficits in adults (Fig 4A). However overexpression of *WT-CaMKII* in adults modestly rescued *FMRFaR^RNAi^* induced flight deficits on day 4, 6 and 8 (Fig 4A, S2A Fig - *THD1;TARGET* controls and S3A Fig – *WT-CaMKII/+;FMRFaR^RNAi^/+* controls). It should be noted that overexpression of *WT-CaMKII* in adult dopaminergic neurons also reduced the duration of flight bouts (S3A Fig; orange bars), suggesting that CaMKII levels need to be tightly regulated in adult dopaminergic neurons for maintenance of flight (see Discussion). Lack of robust rescue by CaMKII of flight deficits in *FMRFaR* knockdown flies may in part arise from an imbalance between levels of FMRFaR and CaMKII in *THD1* neurons. Rescue with the constitutively active form of CaMKII (*CaMKII^T287D^*) was not attempted because expression of *CaMKII^T287D^* in adult *THD1* neurons resulted in strong flight deficits (S3A Fig; maroon bars). Reduced neuronal excitability by overexpression of the mutant transgene, *CaMKII^T287D^* in larval neurons has been observed previously where it elicited behavioral deficits, presumably via modulation of potassium currents (30) (also see discussion). Overall the partial rescue of flight deficits in the *FMRFaR* knockdown animals by expression of a *WT-CaMKII* transgene indicates a genetic interaction between them.

**Fig 4.**
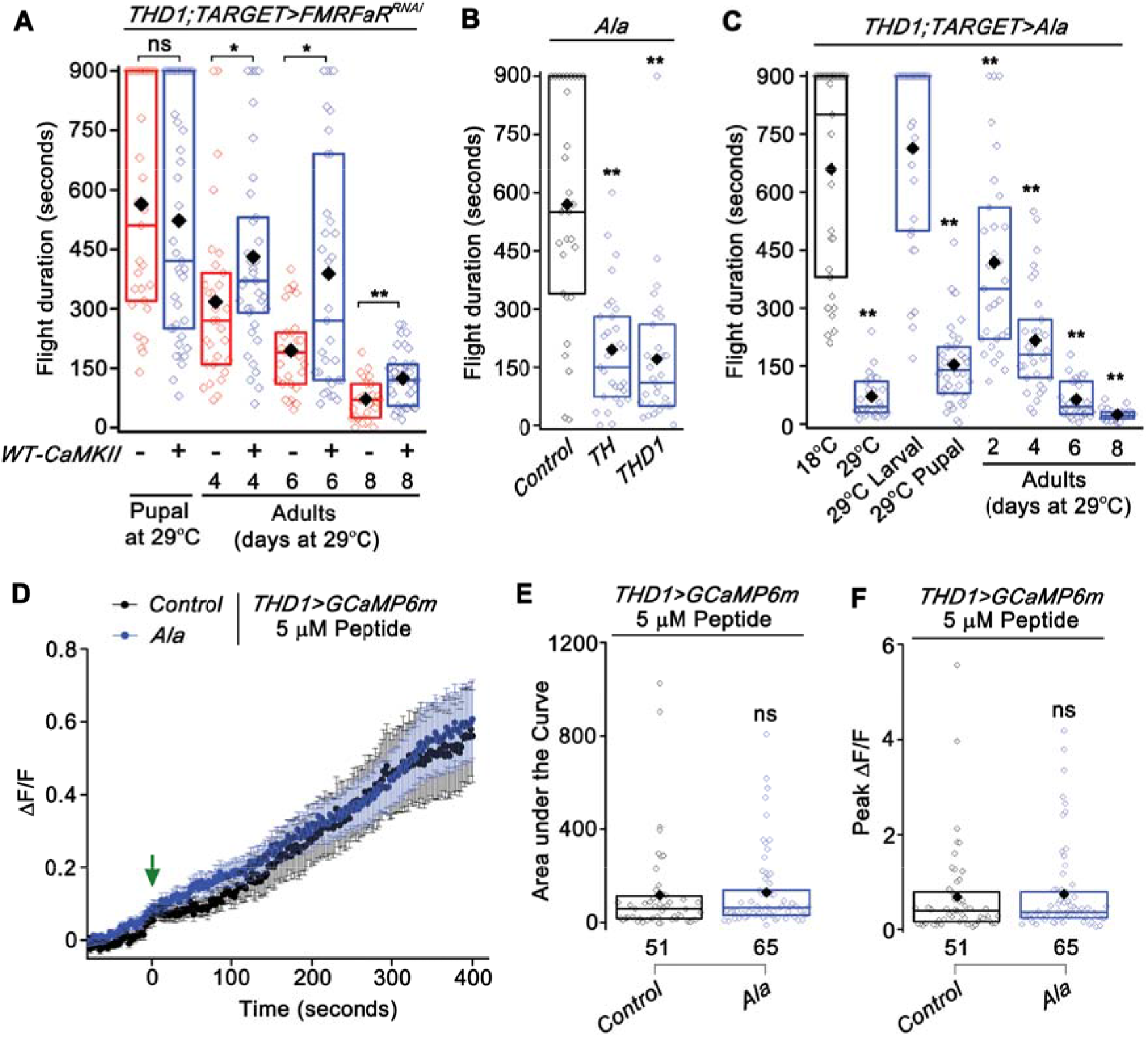
CaMKII function is required in dopaminergic neurons to regulate flight. (A) Flight bout durations observed with overexpression of *WT-CaMKII* in the background of *FMRFaR* knockdown in *THD1* neurons (*THD1;TubGAL80^ts^>WT-CaMKII;FMRFaR^RNAi^*). A significant rescue of the flight defect was observed by expression in the adults, but not in pupae (n≥30, *p<0.05, **p<0.01, Mann-Whitney U-test). (B) Flight durations observed with expression of a genetically encoded CaMKII inhibitor peptide, *Ala*, in dopaminergic neurons as compared to *Ala* control (n≥30, **p<0.01, Mann-Whitney U-test). (C) Flight time of flies with temporal expression of *Ala* in *THD1* neurons using the TARGET system (*THD1;TubGAL80^ts^>Ala*). Significant deficits in flight were observed upon CaMKII inhibition in pupal and adult stages as compared to flies maintained at 18°C throughout (n≥30, **p<0.01, Mann-Whitney U-test). (D) Mean traces (±SEM) of normalized GCaMP6m responses in control (*THD1>GCaMP6m*) and *Ala* expressing *THD1* neurons (*THD1>GCaMP6m;Ala*) upon peptide stimulation (green arrow). (E) Area under the curve and (F) Peak ΔF/F quantified from 0 s to 420 s in (D) were not significantly different between the two genotypes tested (p>0.05; ns - not significant; Mann-Whitney U-test).

To test directly if CaMKII is required in dopaminergic neurons to modulate flight bout durations, we expressed a synthetic peptide inhibitor of CaMKII (*Ala*) (31) in dopaminergic cells of interest. Ala is a peptide analog of the CaMKII autoinhibitory domain that can bind to the catalytic domain of CaMKII, thereby functioning as an exogenous inhibitor. Inhibition of CaMKII in pan-dopaminergic (*TH*) or the subset dopaminergic neurons (*THD1*) resulted in flies with significantly reduced durations of flight bouts (Fig 4B). Knockdown of *CaMKII* with a *CaMKII^RNAi^* also elicited a significant reduction of flight bout durations (S3B Fig). Inhibition of CaMKII activity by the *Ala* peptide in *THD1* neurons did not affect their general locomotor ability, indicating that CaMKII function in *THD1* neurons is likely to be flight specific (S3C Fig). Additionally, TARGET experiments demonstrated that *THD1*-driven expression of *Ala* specifically in either pupal or adult stages significantly compromised the duration of flight bouts, as compared to genetic controls (Fig 4C, S2A Fig - *THD1;TARGET* controls, S3D Fig - *Ala/+* controls). Given that CaMKII is involved in several developmental processes, for example, axon terminal growth (32), a pupal requirement for CaMKII is not surprising. Maximal reduction in flight bout duration was observed in flies wherein CaMKII was inhibited by *Ala* peptide expression in *THD1* cells for 8 days as adults (Fig 4C).

These data support a role for CaMKII activity in the *THD1* subset of dopaminergic neurons in the specific context of maintaining flight bout durations. Similar to *FMRFaR* knockdown, CaMKII inhibition with *Ala* in adult neurons did not affect levels of *TH* mRNA (S3E Fig). Thus, unlike previous findings where reduced Ca^2+^ signaling in pupal dopaminergic neurons resulted in reduced expression of the gene encoding TH (*ple* or *TH*) (6, 7), FMRFaR and CaMKII in adult dopaminergic neurons appear to modulate flight by an alternate cellular mechanism.

To test if CaMKII activity affects FMRFaR induced Ca^2+^ release, *THD1* neurons expressing *Ala* peptide were stimulated with the FMRFa peptide. FMRFa-stimulated cellular calcium responses were not affected by *Ala* expression in *THD1* neurons (Fig 4D, E, F). These data suggest that the function of CaMKII is either downstream or parallel to FMRFaR induced Ca^2+^ release.

The ability of FMRFa to activate CaMKII in *Drosophila* central brain neurons was tested next by immunostaining for pCaMKII, a CaMKII modification that occurs upon prolonged calcium elevation in the cell (30, 33). For technical reasons, this immunostaining was performed on primary neuronal cultures from larvae, wherein we overexpressed the *FMRFaR*^+^ in all *nSybGAL4* positive neurons (34). In addition, the same neurons were marked with *mCD8GFP* to normalize the pCaMKII signal and to account for variability in strength of expression of the overexpressed *FMRFaR^+^*. A significant increase in pCaMKII/GFP staining was observed upon stimulation of larval *nSyb>FMRFaR^+^;mCD8GFP* positive neurons with FMRFa as compared to cells with solvent addition (S3F-H Fig). These data support the idea that FMRFa stimulated Ca^2+^ release activates CaMKII in central brain neurons. Even though a direct test for FMRFa-stimulated CaMKII activation in *THD1* neurons has not been possible so far due to technical reasons, the ability of *WT-CaMKII* to partially rescue the flight deficit in *FMRFaR* knockdown animals (Fig 4A), taken together with these cellular data suggest that CaMKII may function downstream of FMRFaR signaling. Nevertheless, an FMRFaR-independent and parallel role for CaMKII in *THD1* neurons remains possible. The underlying cellular basis for the flight deficits observed upon *FMRFaR* knockdown and *Ala* expression in dopaminergic neurons was investigated next.

### FMRFaR and CaMKII modulate calcium entry and membrane potential in central dopaminergic neurons

To measure neuronal activity of *THD1* neurons, we tested their response to a depolarizing stimulus. *Ex vivo* brain preparations from *THD1* marked dopaminergic neurons were stimulated with 70 mM KCl, a condition known to raise the resting membrane potential and activate voltage-gated calcium channels on the plasma membrane (35). The stimulation was followed by optical measurements of calcium dynamics. Firstly, using the TARGET system, we expressed *GCaMP6m* in *THD1* marked cells, specifically in adults. Animals reared at 29°C for 8 days (also the time corresponding to maximal flight loss upon *FMRFaR* knockdown or *Ala* expression; Fig 3A, 4C), were used in these experiments. Robust Ca^2+^ elevations were observed in *THD1* marked cells from control animals upon addition of KCl (Fig 5A-D; black trace). Ca^2+^ responses from neurons with either *FMRFaR* knockdown or *Ala* expression, were however significantly reduced as compared to controls (Fig 5A-D; red and blue traces). Similar experiments on *ex vivo* brains obtained after *FMRFaR* knockdown in adults for 2 days, exhibit a minimal decrease in Ca^2+^ responses upon depolarization with KCl, in agreement with their weak flight deficits (S4A-C Fig, Fig 3A). These data suggest that FMRFaR and CaMKII help maintain normal cellular responses to changes in membrane excitability in *THD1* neurons. Interestingly, in the *FMRFaR* knockdown condition, most cells exhibit a dampened Ca^2^+ response (Fig 5D, 2^nd^ row - pink arrow), but a few cells responded normally (Fig 5D, 2^nd^ row - yellow arrow). Their normal response may be due to insufficient receptor knockdown. Alternately all *THD1* marked cells might not express the FMRFaR (see discussion).

**Fig 5.**
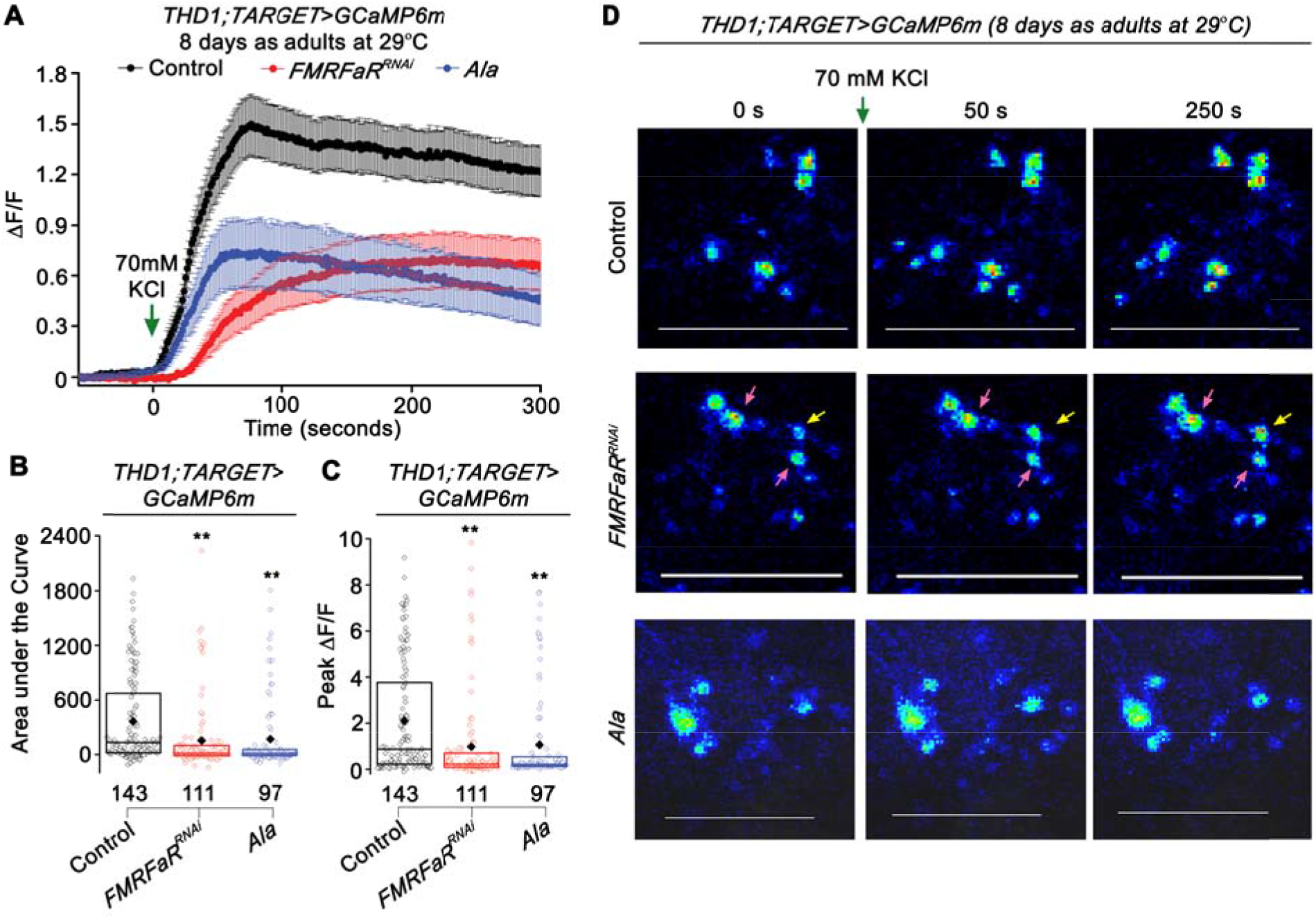
Expression of *FMRFaR^RNAi^* and inhibition of CaMKII in dopaminergic neurons reduces calcium entry after a depolarizing stimulus. (A) Mean trace (±SEM) of normalized GCaMP6m responses observed upon KCl stimulation (green arrow) in 8 day adult neurons in the three indicated genotypes, *THD1;TubGAL80^ts^>GCaMP6m, Control* in black; *THD1;TubGAL80^ts^>GCaMP6m;FMRFaR^RNAi^*, in red and *THD1;TubGAL80^ts^>GCaMP6m;Ala*, in blue. (B) Area under the curve and (C) Peak change in fluorescence quantified from (A). Each bar is compared to control shown in black (**p<0.01; Mann-Whitney U-test). (D) Snap shots of GCaMP6m responses from the same genotypes shown in (A). In the *FMRFaR^RNAi^* knockdown condition (2^nd^ row), the yellow arrow indicates a cell that responded whereas pink arrows indicate cells with no response or a minimal response to the depolarization stimulus. Scale bars represent 50 μm.

### Increased membrane excitability can partially compensate for loss of FMRFaR and CaMKII function

Based on the reduced Ca^2+^ elevation upon depolarization with KCl, observed in *THD1* marked dopaminergic neurons of flies with either *FMRFaR* knockdown or CaMKII inhibition, we hypothesized that such neurons might have reduced membrane excitability. To ascertain if reduced FMRFaR signaling indeed alters the ability of *THD1* neurons to undergo membrane depolarization, a genetically encoded fluorescent voltage indicator, *Arclight* (36), was expressed in dopaminergic neurons of *THD1GAL4*. Stimulation of 8 day old adult brains with KCl showed a robust change in Arclight fluorescence corresponding to neuron depolarization (Fig 6A-D; black trace). As proof of concept, expression of *Kir2.1* in *THD1* cells for 2 days nearly abolished KCl induced membrane depolarization (S5A-C Fig; blue trace). Dopaminergic neurons with *FMRFaR* knockdown for 8 days displayed a significantly reduced ability to depolarize after KCl addition (Fig 6A-D; red trace). As was observed previously (Fig 5D), membrane depolarization induced changes in Arclight fluorescence were attenuated in most *THD1* neurons (Fig 6D, 2^nd^ row; pink arrow), while few responded like controls (Fig 6D, 2^nd^ row; yellow arrow). Similar experiments with adult brains at 2 days exhibit equivalent changes in membrane potential in both control and with *FMRFaR^RNAi^*, in agreement with the weaker flight deficits of 2d old flies with *FMRFaR* knockdown (S5A-C, Fig 3A).

**Fig 6.**
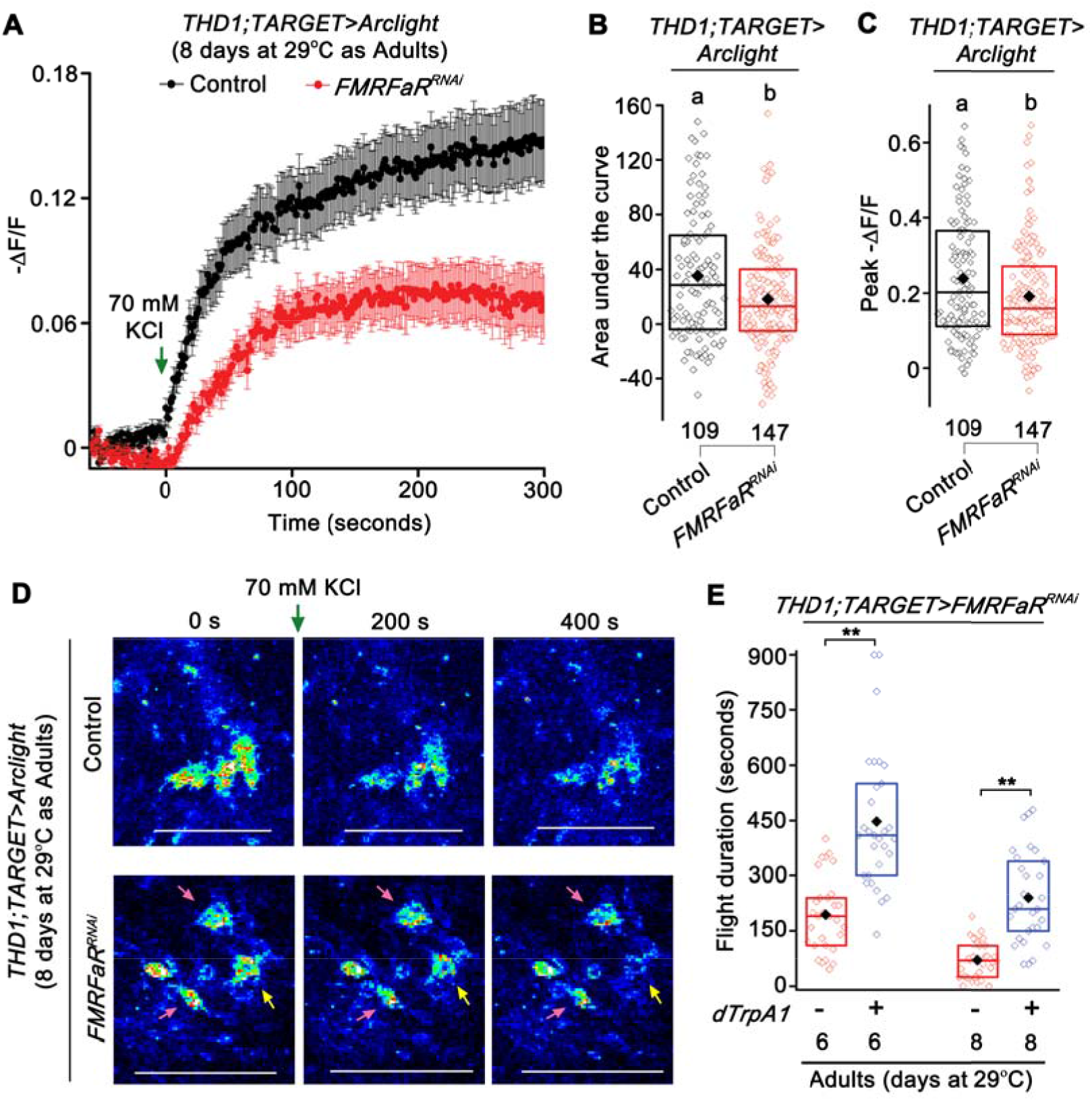
Expression of *FMRFaR^RNAi^* in dopaminergic neurons reduces membrane depolarization. (A) Mean trace (±SEM) of negative change in fluorescence (-ΔF/F) observed upon KCl stimulation of *THD1* neurons expressing a fluorescent-based membrane voltage indicator, *Arclight* in the indicated genotypes, *THD1;TubGAL80^ts^> Arclight, Control* in black; *THD1;TubGAL80^ts^>Arclight;FMRFaR^RNÄl^*, in red. (B) Area under the curve and (C) Peak (-ΔF/F) quantified from (A) were significantly different in the *FMRFaR* knockdown neurons as compared to their genotypic controls. Numbers below each box plot indicate total number of cells imaged from a minimum of 5 independent brains (One-way ANOVA followed by post-hoc Tukey’s test; the same alphabet above each bar represents statistically indistinguishable groups; different alphabet represents p<0.05). (D) Representative images of neuronal depolarization as observed by a decrease in Arclight fluorescence in the same genotypes shown in (A). With *FMRFaR* knockdown (2^nd^ row), few cells responded normally (yellow arrow), whereas most others showed no response or a minimal response to the depolarization stimulus (pink arrow). Scale bars represent 50 μm. (E) Rescue of flight bout durations by expression of *dTrpA1* in the genetic background of *FMRFaR^RNAi^* in 6 and 8 day old adult *THD1* marked neurons (*THD1;TubGAL80^ts^>TrpA1;FMRFaR^RNAi^* compared to *THD1;TubGAL80^ts^>FMRFaR^RNAi^;* n≥30, **p<0.01, Mann-Whitney U-test).

Because knockdown of *FMRFaR* reduced KCl induced Ca^2+^ entry in dopaminergic neurons (Fig 5A-D), attenuated membrane depolarization (Fig 6A-D) and correlated with shortened flight bout durations (Fig 3A), we hypothesized that FMRFaR function is likely required to maintain optimal excitability in these neurons. Consequently, increasing neuronal excitability might compensate for reduced FMRFaR function and thereby rescue the flight deficits observed in *FMRFaR* knockdown flies. To test this idea, we introduced a temperature sensitive cation channel, *dTrpA1* (37) in flies with adult specific *FMRFaR* knockdown. Activation of the dTrpA1 calcium channel can compensate in part for reduced Ca^2+^ entry through voltage-gated channels (15, 38). Indeed, expression of *dTrpAl* significantly improved the maintenance of flight bouts in 6 and 8 day old *FMRFaR* knockdown adults (Fig 6E, S2A Fig - *THD1;TARGET* controls and S5D Fig - *TrpA1/+;FMRFaR^RNAi^/+* controls). Likewise, expression of *NaChBac* (a bacterial sodium channel) (39), moderately alleviated the near loss of flight observed with Ala-dependent CaMKII inhibition in 6 and 8 day old adults (S5E, F Fig). Expression of another UAS element (*GCaMP6m*) in the background of *FMRFaR* knockdown or *Ala* expression did not rescue the loss of flight (S5F Fig; green bar, as compared to Fig 3A; orange bar – 8 days as Adults at 29°C; p>0.05; Mann-Whitney U-test). Expression of just the *TrpA1* or the *NaChBac* transgenes in adult dopaminergic neurons also affected flight bout durations, supporting the idea that optimal activity of ion channels in *THD1* marked adult dopaminergic neurons is required for their normal function (S5D, F Fig). Taken together, these data demonstrate that FMRFaR helps maintain optimal excitability of flight bout extending central dopaminergic neurons.

### Membrane excitability and synaptic vesicle recycling in dopaminergic neurons is essential for sustained flight

To directly test if changes in membrane excitability of *THD1* neurons, affects their ability to modulate flight bout durations, we expressed transgenes that encode ion channel specific tethered-toxins in adult *THD1* neurons. Expression of such toxins, in *Drosophila* neurons, inhibits voltage-gated ion channels thereby altering neuronal excitability and behavior (40). Expression of *PLTXII*, a toxin that targets the presynaptic calcium channel, *cacophony*, in adult *THD1* neurons significantly impaired maintenance of flight bouts from as early as 2 days of transgene expression (Fig 7A, S6A Fig). By 6 days, just 20% of the flies survived *PLTXII* expression and hence flight measurements were recorded only up to 4 days at 29°C (Fig 7A, S6A Fig). Similarly, blocking potassium channels by expression of *κ-ACTX-Hv1c*, in adult dopaminergic neurons also compromised the duration of flight bouts to a significant extent (Fig 7B, S6A Fig). Approximately 60% lethality was observed with *κ-ACTX-Hv1c* expression in adult *THD1* neurons for 6 days. It is not surprising that we observe lethality with the *PLTXII* and *κ-ACTX-Hv1c* transgenes as pan-neuronal expression of each of these toxins has previously been reported to result in lethality (40). Next, adult specific expression of *δ-ACTX-Hv1a*, a toxin that inhibits inactivation of voltage-gated sodium channels (encoded by the gene *para*), in *THD1* neurons also reduced flight bout duration in adult flies after 4 days of transgene expression (Fig 7C, S6A Fig). Furthermore, by immunostaining for endogenous TH, we confirmed that dopaminergic neurons were intact under these conditions (S6B Fig). Taken together, these data suggest that the membrane excitability of flight modulating *THD1GAL4* marked central dopaminergic neurons must be tightly controlled for maintenance of long flight bouts.

**Fig 7.**
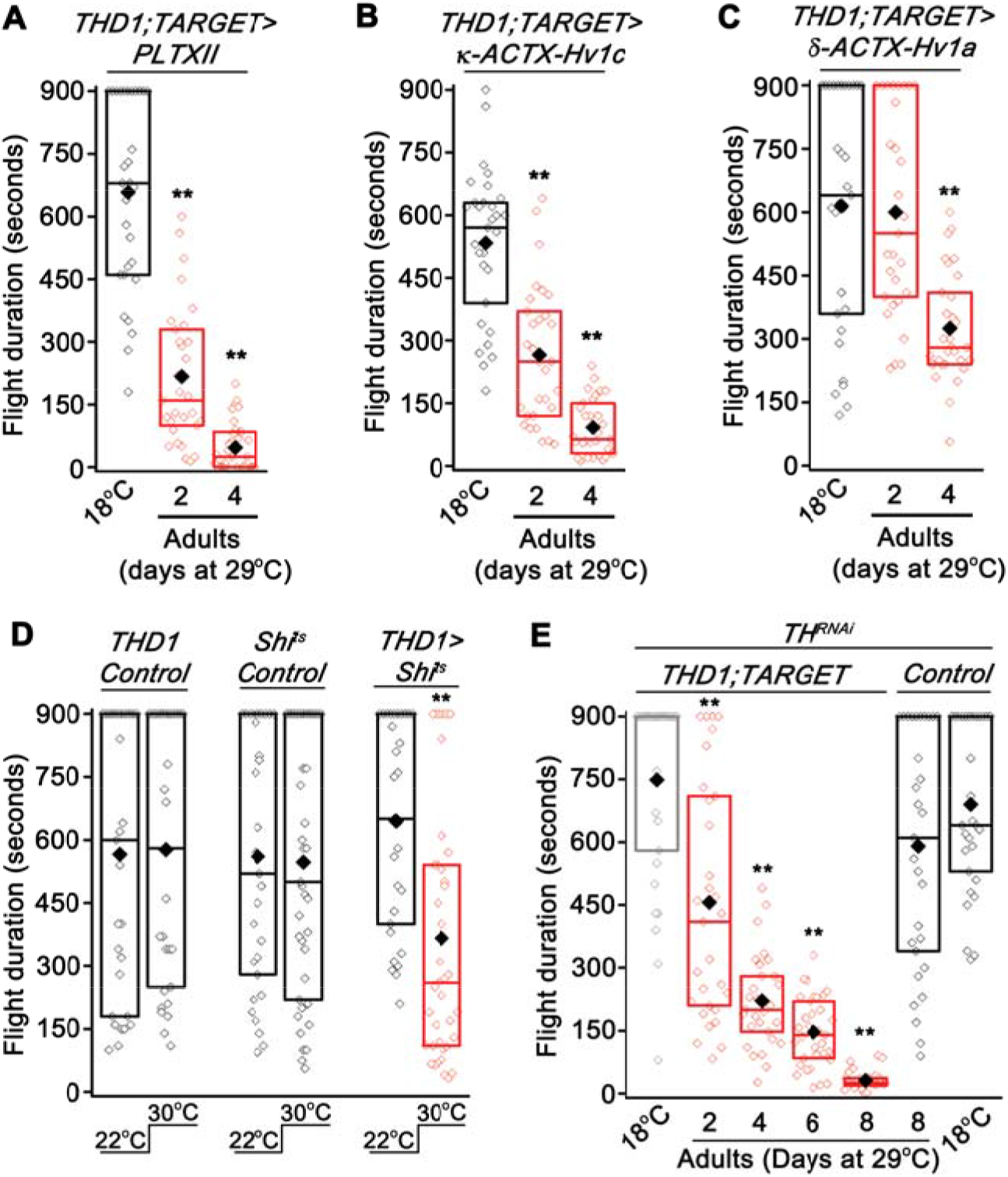
Dopaminergic neurons require optimal membrane excitability and synaptic vesicle recycling for sustained flight. (A) Flight bout durations were significantly reduced upon adult specific expression of the Ca^2+^ channel inhibitor, *PLTXII (THD1;TubGAL80^ts^>PLTXII;* n≥30, **p<0.01, Mann-Whitney U-test). (B) Expression of a K+ channel inhibitor, *κ-ACTX-Hv1c*, in adult dopaminergic neurons reduces flight bout durations (*THD1;TubGAL80^ts^>κ-ACTX-Hv1c;* n≥30, **p<0.01, Mann-Whitney U-test). (C) Adult dopaminergic neuron specific expression of *δ-ACTX-Hv1a*, a toxin that targets Na+ channels, significantly reduces flight bout durations (*THD1;TubGAL80^ts^>δ-ACTX-Hv1a;* n≥30, **p<0.01, Mann-Whitney U-test). In all the above cases (A, B, C), comparisons have been made with the corresponding 18°C control (black). (D) Acute expression of a temperature sensitive dynamin mutant (*Shi*^ts^) in adult dopaminergic neurons reduced flight bout durations when tested at 30°C as compared to a control tested at 22°C (*THD1>Shi^ts^* at 22°C compared to 30°C; n≥30, **p<0.01, Mann-Whitney U-test). *THD1/+* and *Shi^ts^/+* controls in the two temperature conditions are also shown. (E) Adult specific knockdown of the dopamine synthesizing enzyme, Tyrosine Hydroxylase (*TH*) by expressing an RNAi (*TH^RNAi^*) in dopaminergic neurons significantly reduces flight duration as compared to the corresponding 18°C controls (*THD1;TubGAL80^ts^>TH^RNAi^;* n≥30, **p<0.01, Mann-Whitney U-test).

We hypothesized that changes in membrane excitability are likely to affect the release of dopamine containing synaptic vesicles from *THD1* neurons. A temperature-sensitive mutant transgene for dynamin (*UAS-Shibire^ts^* or *Shi^ts^*) (41) was used to block synaptic vesicle recycling by shifting flies transiently to the non-permissive temperature of 30 C. Indeed, when synaptic vesicle recycling was inhibited in *THD1* neurons of adult flies, reduced flight bout durations were observed as compared with controls (Fig 7D). Flies of the same genotype (*THD1>Shi^ts^*) maintained at 22°C throughout showed normal flight bout durations (Fig 7D). Thus acute manipulation of synaptic function of adult *THD1* neurons affects the maintenance of flight bout duration in the tethered flight paradigm. Finally, we tested requirement for the neurotransmitter, dopamine in *THD1* neurons, for modulating the duration of adult flight bouts. Adult-specific expression of an RNAi transgene for the dopamine-synthesizing enzyme Tyrosine Hydroxylase (*TH^RNAi^;* (42)) in *THD1* neurons significantly impaired the maintenance of longer flight bouts from as early as 2 days of *TH^RNAi^* expression (Fig 7E), suggesting that dopamine release from synapses of *THD1* neurons is required for maintaining longer flight bouts.

## Discussion

In this study we demonstrate that the FMRFaR and CaMKII signaling in specific central dopaminergic neurons of the adult *Drosophila* brain helps maintain optimal membrane excitability (Fig 5A-D, 6A-D). Previous work by our group demonstrated that loss of IP_3_R mediated Ca^2+^ signaling in central dopaminergic neurons during pupal stages similarly affects flight circuit function in adults (6, 7). In parallel, neuropeptidergic modulation of flight by the Pdf receptor was investigated, but the exact class of neurons that required this receptor or the downstream effectors remained elusive (14). For the first time, we describe here a neuropeptidergic signaling mechanism in adult brain neurons that is required for sustained flight (S6C Fig). Our data support a model where FMRFa-modulated activity in central dopaminergic neurons belonging to the PPL1 and PPM3 clusters extends the duration of flight bouts, a behavior that is likely to have significant effects on optimal sourcing of nutrients, finding sites for egg laying and identifying mates in the wild.

The role of neuropeptide signaling in modulating animal behavior is well documented (2, 4). Indeed, rescue of flight deficits by overexpression of the cation channel *TrpA1* in dopaminergic neurons of *FMRFaR* knockdown flies directly addresses the importance of peptide driven neuronal excitability, for flight behavior (Fig 6E). Reduced cellular responses to a depolarizing stimulus observed upon *FMRFaR* knockdown in *THD1* neurons may arise from functional modification, possibly by CaMKII, of plasma membrane resident ion channels, or their expression levels (Fig 5A-D, 6A-D). However, our data do not exclude other kinases or modifiers as functioning downstream of the FMRFaR. We predict that FMRFaR in dopaminergic neurons is a key neuromodulator of membrane excitability which stimulates release of dopamine containing synaptic vesicles. In fact, stimulation of synaptic transmission downstream of FMRFaR activation has been described previously in *Drosophila* larval neuromuscular junctions (13). Though we have not tested synaptic vesicle release directly upon FMRFaR stimulation, our data support an essential role for synaptic transmission and dopamine synthesis in adult dopaminergic neurons for maintenance of flight bout durations (Fig 7D, E). Identification of the precise ion channels that are affected by FMRFaR signaling, leading to changes in membrane excitability and very likely synaptic transmission in flight modulating dopaminergic neurons, needs further investigation.

Multiple FMRFa peptides exist in *Drosophila* (26, 27). Cells positive for FMRFa peptides have been characterized in the *Drosophila* central nervous system using antisera specific to some of the FMRFa peptides (43–46). Immunostaining of adult brains, especially against the peptide, DPKQDFMRFa (also used in this study) have revealed extensive neuronal varicosities in the anterior and lateral regions of the protocerebrum (45, 46). It is hence conceivable that FMRFa released through these projections activates the FMRFaR on the anatomically proximal PPL1 and PPM3 clusters of dopaminergic neurons. The PPL1 neurons are known to innervate the superior protocerebrum, a region considered to function at the interface of olfactory input and motor output modules of the brain (47). Thus activation of the FMRFaR on dopaminergic neurons might stimulate dopamine release, which could then reinforce a dopaminergic circuit, required for sustained flight. Furthermore, co-expression of the dopamine synthesizing enzyme Tyrosine Hydroxylase (TH) in a few neurons marked by an *FMRFaGAL4* (48), suggests the existence of an autocrine signaling mechanism within the flight modulating dopaminergic neurons. This idea is supported by recent studies of single cell sequencing of *Drosophila* neurons where FMRFa transcripts were seen in a small number of central brain dopaminergic neurons (49, 50). Interestingly, a recent report from *Manduca* demonstrated that an FMRFa positive neuron lies at the center of a putative sensory-motor circuit for integration of olfactory stimuli with wing movements during flight (51). The natural context in which FMRFa release is triggered for modulation of flight in *Drosophila* remains to be identified. Neuromodulatory signals from other receptors in central dopaminergic neurons is also a possibility and would widen the scope of sensory inputs received by these cells for integration with flight behavior.

FMRFaR activation by the peptide DPKQDFMRFa stimulates intracellular Ca^2+^ release through the IP_3_R (Fig 2D-F) (13) and enhances synaptic transmission at the larval neuromuscular junction and thereby modulates muscle contraction (13, 29, 46). Previous work from our lab has demonstrated that in *Drosophila* neurons, subsequent to intracellular Ca^2+^ release, cytosolic calcium is further elevated by store-operated Ca^2+^ entry (SOCE) through dOrai (52). Thus, we predict that the rise in Ca^2+^ we see upon FMRFa stimulation of *THD1* neurons could possibly be a combination of both IP_3_R mediated Ca^2+^ release and SOCE. Rescue of flight deficits in the *FMRFaR* knockdown flies by an *itpr*^+^ transgene (Fig 3A) and loss of peptide induced Ca^2+^ rise in an *itpr* knockdown background (Fig 2D-F) support the 2^+^ idea that IP_3_R-mediated Ca^2+^ release is likely downstream of FMRFaR activation. Activation of overexpressed IP_3_Rs in the *FMRFaR* knockdown condition might occur through alternate receptors. It is also possible that overexpression of the IP_3_R allows for more ER-Ca^2+^ release from the residual FMRFaRs after RNAi knockdown. Thus, in the context of flight as well, FMRFaR signaling in dopaminergic neurons appears to function upstream of IP_3_R-mediated Ca^2+^ release (Fig 2D-F, 3A). The adult requirement of FMRFaR on dopaminergic neurons supports a primary role for this receptor in mature neurons for modulation of flight, but not as much for maturation of the flight circuit in pupal stages (Fig 3A). In contrast, the dFrizzled2 receptor, which also functions upstream of the IP_3_R in central dopaminergic neurons, was shown to be required exclusively during the pupal stages, albeit in a different dopaminergic cluster of the brain (6). These observations support the existence of specific receptors that stimulate IP_3_-mediated Ca^+2^ release and function either during flight circuit maturation in pupae or in a neuromodulatory role in adults, to influence flight circuit dynamics. A range of flight deficits in IP_3_R mutants with differing allelic strength support this idea (20).

As described previously by other groups in larval motor neurons and in larval neuromuscular junctions (13, 29), our genetic and cellular data also suggests that CaMKII might be a downstream effector of FMRFaR signaling in neurons (Fig 4A, S3F-H Fig). Although we have shown increased pCaMKII immunostaining in larval CNS neurons upon FMRFa stimulation (S3F-H Fig), direct activation of either CaMKII or other Ca^2+^ dependent kinases by FMRFa in adult *THD1* neurons, remains to be tested rigorously. CaMKII, thus far has been known to contribute to synaptic plasticity in the context of learning and memory (53, 54). Our data, for the first time, demonstrate that CaMKII is required in dopaminergic neurons for maintenance of *Drosophila* flight over periods of several minutes, possibly by modulating neuronal firing during flight. Multiple mechanisms for CaMKII dependent modulation of membrane excitability have been described in *Drosophila* such as phosphorylation of the potassium channel, Eag, leading to an increase in the Eag current (55). In another study, CaMKII dependent phosphorylation of the Ca^2+^ activated potassium channel binding protein, Slob was shown to favor its binding to 14-3-3, that eventually altered the voltage sensitivity of slowpoke channels (56). In both cases, CaMKII acted as a negative regulator of neuronal excitability, and support our data demonstrating flight deficits observed upon overexpression of *WT-CaMKII* and *CaMKII^T287D^* (S3A Fig). However, the more interesting and compelling explanation for our data demonstrating reduced KCl-induced depolarization upon CaMKII inhibition by *Ala* comes from *in vitro* work that has described a role for CaMKII in decelerating the inactivation of voltage sensitive calcium channels, thereby enhancing transmitter release (57). This was however shown to be independent of its kinase activity. More recently, there is evidence to suggest CaMKII-dependent activation of transcription factors in *Drosophila* neurons (58, 59). Hence, the possibility of transcriptional regulation of voltage-gated membrane channel genes by CaMKII cannot be excluded and needs further investigation.

## Materials and Methods

### Fly rearing and stocks

*Drosophila* strains used in this study were maintained on cornmeal media, supplemented with yeast. Flies were reared at 25°C, unless otherwise mentioned. WT strain of *Drosophila* used was Canton-S. The fly lines *nSybGAL4* (BL51635), *UASFMRFaR^RNAi^* (BL25858), *UAScas9* (BL54593), *UASdTrpA1* (BL26263), *UASNaChBac* (BL9468), *UASKir2.1* (BL6595), *UASGCaMP6m* (BL42748), *UASArclight* (BL51057), *UASmCD8GFP* (BL5130), *UASAla* (BL29666), *UASWT-CaMKII* (BL29662), *UASCaMKII^T287D^* (BL29664), *UASTH^RNAi^* (BL25796), *UASDicer* (BL24648) were obtained from Bloomington *Drosophila* Stock Centre (BDSC). The *UASitpr^RNAi^* (1063-R2) strain was from National Institute of Genetics (NIG) and the *UASCaMKII^RNAi^* (v38930) was from Vienna Drosophila Resource Center (VDRC). A strain with two copies of *TubGAL80^ts^* on the second chromosome was generated by Albert Chiang, NCBS, Bangalore, India, and has been used for all TARGET experiments. The dopaminergic GAL4 driver, *THGAL4* (21) and the *THGAL80* strain (23) were kindly provided by Serge Birman (CNRS, ESPCI Paris Tech, France). All the TH subset GAL4s (*THD1, THD’, THC1, THC’*) were a gift from Mark N Wu (Johns Hopkins University, Baltimore) (22). The *UASeGFP* transgene was provided by Michael Rosbash (Brandeis University, Waltham, MA). The *UASShi^ts^* strain was obtained from Toshihiro Kitamoto (University of Iowa, Carver College of Medicine, Iowa City). The *UAS-toxin* fly strains have been described earlier (40) and were obtained from Brian McCabe (EPFL Brain Mind Institute, Lausanne, Switzerland). The generation and use of *UASitpr*^+^ transgene has been described earlier (15, 60). The *UASFMRFaR^+^* strain was generated in the lab (unpublished). Standard fly genetics were followed for making strains and recombinants.

Temperature shift experiments were performed as described below. Briefly, larvae, pupae or adults were maintained at 18°C and transferred to 29°C only at the stage when the *UAS-transgene* needed to be expressed. Firstly, as a control, flies of the genotype *THD1;TARGET>UAS-transgene* were maintained throughout at 18°C, a condition where the *UAS-transgene* expression is suppressed (labeled as ‘18°C’ in figures). To observe maximum effect, flies of the same genotype were grown at 29°C throughout development and as adults (labeled ‘29°C’). For larval specific knockdown, animals of the desired genotypes developed at 29°C from embryos to the wandering larval stage, following which they were maintained at 18°C till the time when they were tested for flight (labeled as ‘29°C Larval’). Likewise, for pupal specific knockdown, wandering third instar larvae were shifted from 18°C to 29°C. Pupae developed at 29°C and were transferred to 18°C, soon after eclosion (labeled as ‘29°C Pupal’). In case of adult specific knockdown, larval and pupal development continued at 18°C. Following this, adult flies were maintained at 18°C for two days post eclosion and then transferred to 29°C and used for experiments at 1, 2, 4, 6 or 8 days after transfer (labeled as ‘29°C Adult’).

### Generation of a knockout for *FMRFaR* using the CRISPR-Cas9 system

To generate a *FMRFaR* null allele, we used the CRISPR-Cas9 methodology (61 – 65). Two guide RNAs (gRNAs) were designed, at the 5’ and 3’ends of the gene at the following sequences: (sg1-GGGAGCCATGAGTGGTACAGCGG, sg4-GATCTCTGCATTTCGCGGGCGGG) so as to delete ~1.5 kb from the coding region of the 1.6 kb gene. A gRNA dual transgenic fly (*FMRFaR^dual^*) was made first (24). *FMRFaR^dual^* transgenic males were mated with *Act5c-Cas9* virgins (66). From the F1 progeny obtained, 16 were screened by PCR for the deletion. Because all flies tested were positive for the deletion, 4 F1 flies were individually crossed to balancers and 30 F2 progeny were screened for the deletion. Progeny, that were positive for deletion and negative for presence of dual gRNA (total 6 F2s), were maintained as stable lines. The deletion was confirmed by sequencing of the DNA from the *FMRFaR* knockout flies. The following primers were used for confirmation of deletion - 5’F: GACATAGTCATCAGGTGCTC, 5’R: TGCACCTCCGTGTGGTTAAG, 3’F: GAACAACGGCGATGGAACTC, 3’R: GGTGCTCTAAGTCAACCCCT. For tissue specific deletions of *FMRFaR* gene, a fly strain containing *FMRFaR^dual^* and *UAScas9* was made and subsequently mated with GAL4 strains of interest.

### Single flight Assay

Single flight tests were performed on adult flies 3-5 days post eclosion, unless specified differently. Flies were anaesthetized on ice for 2-3 minutes and then tethered using nail polish applied to the end of a thin metal wire. The tether was glued onto the region between the neck and thorax of adult flies. A gentle mouth blown air-puff was delivered to test for flight response. Flight time in tethered condition was recorded for a maximum of 15 minutes. In three independent batches of 10, a minimum of 30 flies were tested for each genotype. All control genotypes tested were obtained by crossing the GAL4 or UAS strain of interest to the wild-type strain, *Canton-S*. Raw data was plotted as box and whisker plots using Origin (OriginLab, Northampton, MA) software. Each bar represents 25^th^ to 75^th^ percentile. The solid diamond and horizontal line within the bars indicate mean and median, respectively. Individual values are shown as open diamonds.

### Locomotor Assay

Locomotor activity was measured by modifying a previously published protocol (67). Briefly, locomotor activity of 4-6 day old singly housed virgin males was tested in a circular chamber of diameter 4 cm. The chamber was placed on a sheet containing a specific pattern. Single flies were aspirated into each chamber and allowed to acclimatize for 5 minutes. Each fly was monitored separately for the number of times it crossed each line in the pattern, over a period of 10 seconds. For every experiment, six single flies were tested sequentially for the span of 10 seconds. This was repeated 5 times, i.e. we measured locomotor activity of every single fly five times in blocks of 10 seconds, so as to randomize activity measurements during the experimental time. Total locomotor activity is represented as the sum total of the number of lines crossed by each fly over the experimental duration of 50 seconds (plotted as Locomotor Activity Units). A minimum of 25 flies were tested from each genotype.

### *Ex vivo* live imaging of adult brains

Adult brains were dissected in adult haemolymph-like solution (AHL) (68) and embedded in ~10 μl of 1% low melt agarose (Invitrogen) with anterior side facing upwards. Brains were then bathed in 100 μl AHL until imaging. Images were obtained on an Olympus FV1000 inverted microscope (Olympus Corp., Japan) in time lapse mode using the 20X objective. Both GCaMP6m and Arclight signals were captured using the 488 nm excitation laser line. Fifty frames of basal activity were recorded, followed by stimulation with either 5 μM Peptide (DPKQDFMRFa, NeoBioLab, Massachusetts, USA) or 70 mM KCl. A minimum of 5 brain explants were imaged for every experimental condition.

Raw fluorescence data for regions of interest were extracted using the Time Series Analyser plugin in Image J 1.47v. Change in fluorescence, ΔF/F was calculated using the following formula for each time (t): ΔF/F = (F_t_-F_0_)/F_0_, where F_0_ is the average basal fluorescence of the first 10 frames. Area under the curve and Peak ΔF/F was calculated using Microsoft Excel considering all frames post-stimulation. Values were plotted as box plots.

### Sorting of adult neurons by FACS

Twenty adult CNS (*THGAL4>eGFP*) were dissected in cold Schneider’s medium (#21720-024, Life Technologies) followed by ~45 minutes incubation in an enzymatic solution containing 0.75 μg/μl of Collagenase (Sigma-Aldrich) and 0.40 μg/μl Dispase (Sigma-Aldrich). Brain lysates were then spun at 3000 rpm for 3 minutes. The supernatant was discarded and the pellet was resuspended in cold Schneider’s media. Single cell suspensions were obtained by gentle tituration and then passed through a 40 μm mesh filter. Fluorescence-activated cell sorting (FACS) of samples was carried out as described earlier (7). A minimum of 4 biological replicates were sorted and ~10,000 GFP +ve and GFP -ve cells were collected separately in TRIzol Reagent (Life Technologies) for RNA isolation.

### RNA isolation and qPCR

Total RNA was extracted from sorted cells or dissected adult CNS using TRIzol RNA extraction protocol (Ambion, ThermoFischer Scientific). In case of dissected CNS, a minimum of three biological replicates, each containing 5 brains, was used per genotype. Approximately 500 ng of RNA was then used for DNAse treatment and first strand cDNA synthesis. Quantitative real-time PCR was performed on ABI7500 fast machine (Applied Biosystems) using Kapa SYBR FAST qCR Master mix (KAPA Biosystems, Wilmington, MA). *rp49* was used as an internal control to quantify relative levels of the target transcripts. qPCR values are represented as normalized fold change of mRNA transcripts of the indicated genes. In case of sorted cells, normalized mRNA levels of genes in GFP -ve cells have been plotted relative to the GFP +ve population. Every qPCR run was followed by a melt curve analysis to confirm primer specificity. The following formula was used to calculate fold change:

Fold change = 2^-ΔΔCt^, where, ΔΔCt = (Ct(_target gene_)-Ct(_rp49_))_Experimental_ - (Ct(_target gene_)- Ct(_rp49_))_Control_

Sequences of primers used are as follows:

*rp49:*

Forward- CGGATCGATATGCTAAGCTGT

Reverse- GCGCTTGTTCGATCCGTA

*FMRFaR:*

Forward- GTGCGAAAGTTACCCGTCG

Reverse- TAATCGTAGTCCGTGGGCG

*TH (ple):*

Forward- GTTGCAGCAGCCCAAAAGAAC

Reverse- GAGACCGTAATCATTTGCCTTGC

*dSTIM*:

Forward- GAAGCAATGGATGTGGTTCTG

Reverse- CCGAGTTCGATGAACTGAGAG

*dOrai:*

Forward- GAGATAGCCATCCTGTGCTGG

Reverse- CGGATGCCCGAGACTGTC

### Immunohistochemistry

Immunohistochemistry was performed on dissected adult CNS as described earlier (7). Briefly, brains were dissected in 1x PBS, followed by fixation in 4% paraformaldehyde and blocking in 0.2% phosphate buffer, 0.2% Triton-X 100 and 5% normal goat serum. Overnight incubation with primary antibodies was followed by 2 hours of incubation with corresponding fluorescent secondary antibodies. The primary antibodies used were: mouse anti TH (1:50, #22941, ImmunoStar) and rabbit anti GFP (1:10,000, A6455, Life Technologies). The following were the fluorescent secondary antibodies used at 1:400: anti-mouse Alexa Fluor 568 (#A11004, Life Technologies) and anti-rabbit Alexa Fluor 488 (#A11008, Life Technologies). Images were obtained as confocal stacks of 1 μm thickness using 20X objective on Olympus FV1000 Confocal microscope.

### Primary neuronal culture and immunostaining

Primary neuronal cultures and immunostaining of neurons was carried out as described previously (34). Briefly, third instar larval CNS were dissected in Schneider’s medium containing 50 μg/ml Streptomycin (Invitrogen), 10 μg/ml Amphotericin B (Invitrogen) and 50U/ml Penicillin (Invitrogen). Following this, brains were subjected to 40 minutes of enzyme treatment in 0.75 μg/ml Collagenase and 0.40 μg/ml Dispase. Neurons were dissociated, spun down and plated in growth medium, DMEM/F12-1065 (Gibco) supplemented with the antibiotics mentioned previously and 20mM HEPES. All cell culture reagents were procured from Sigma-Aldrich, unless specified differently. Fourteen-sixteen hours old cultures were washed twice with hemolymph-like saline, HL_3_ and incubated with either the FMRFa peptide or a solvent control for 20 minutes. Following this, cells were fixed in 3.5% paraformaldehyde for 20 minutes at room temperature and then washed in wash buffer (1/10^th^ dilution of blocking buffer). Cells were then permeabilized for a total of half an hour (10 minutes * 3 times), blocked in blocking buffer (5% BSA, 0.5% Triton X in PBS) for 1 hour, at room temperature and subsequently incubated overnight in the primary antibody diluted in wash buffer (mouse anti pCaMKII-22B1; 1:100; #sc-32289, Santa Cruz). The next day, cells were washed thrice in wash buffer (for 10 minutes each) and incubated with anti-mouse Alexa Fluor 568 (#A11004, Life Technologies) secondary antibody for half an hour at room temperature. Control dishes for the secondary antibody were treated as the experimental dishes and were incubated with the fluorescent secondary, but not the primary antibody. Following another set of three washes (15 minutes each) in wash buffer, cells were covered in 60% glycerol before imaging. An inverted Olympus FV3000 confocal microscope with a 60X oil objective (1.42 NA), pinhole size 150 μm was used to image the cells. GFP and pCaMKII signals were captured with 488 and 561 laser lines respectively. The same settings were used to capture images from all experimental conditions, on any particular day of imaging. The entire cell volumes of cells were captured as optical slices of 0.60 μm thickness. Images were analyzed in Image J as described above. After applying background subtraction, fluorescence values of pCaMKII were divided over GFP values to obtain the ratios shown in S3G Fig.

### Data representation and analysis

Origin (OriginLab, Northampton, MA) software was used for plotting raw data and calculation of statistical significance. Raw data were tested for normality. A Student t-test or an ANOVA statistical test was performed on normally distributed data. Non-normal data were tested for statistical significance using the Mann-Whitney U-test.

## Acknowledgments

We thank Prof. Jean-Francois Ferveur for helping us set up the locomotor assay, Trayambak Pathak for standardising protocols for the Arclight experiments and Shlesha Richhariya for help with FACS sorting. We thank the Fly Facility at NCBS for generating transgenic fly lines and Dr. Krishnamurthy and NCBS Central Imaging and Flow Facility for help with confocal imaging and FACS. We also thank the Bloomington *Drosophila* Stock Center (National Institutes of Health P40OD018537) for several fly stocks used in this study.

---

**S1 Fig.**
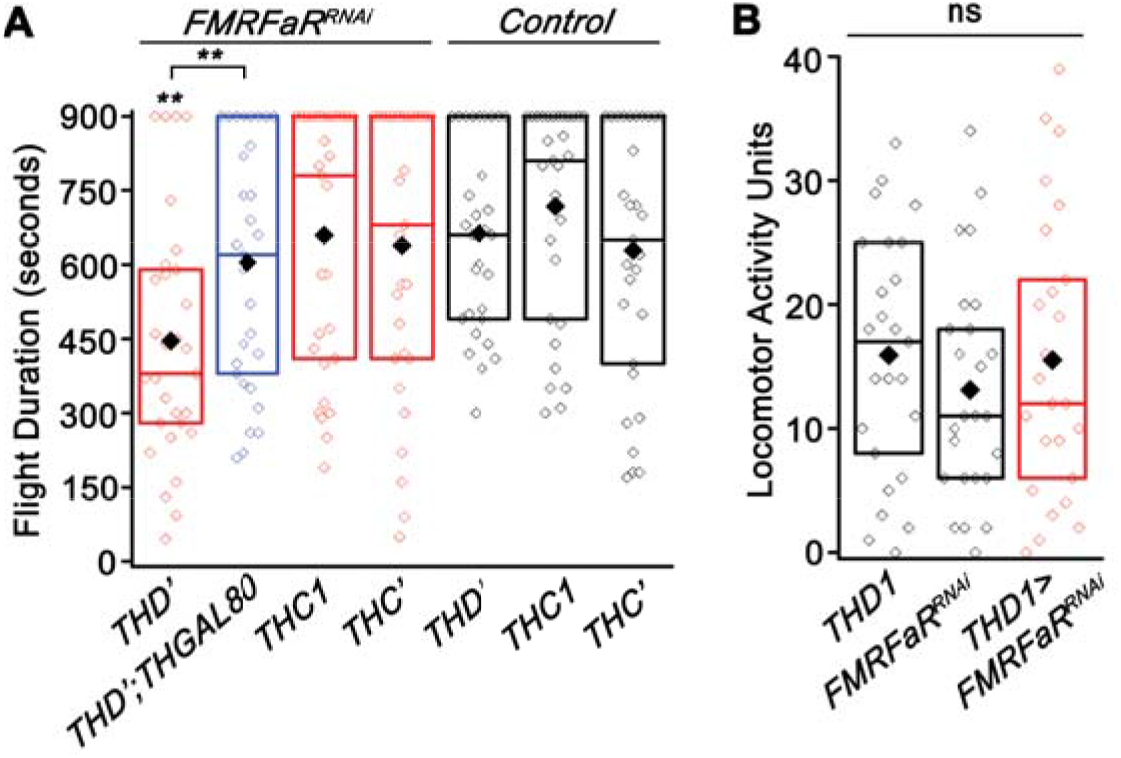
Flight duration and locomotor activity measurements with *FMRFaR* knockdown in subset dopaminergic neurons. (A) Flight durations observed with knockdown of *FMRFaR* in a few TH subset domains, namely, *THD ‘, THC1* and *THC’* as compared to their genotypic controls (*THD’/+, THC1/+, THC’/+* in black and *FMRFaR^RNAi^/+* shown in Fig 1A; n≥30, **p<0.01, Mann-Whitney U-test). The ‘+’ in all control genotypes denotes the wild-type, *Canton-S*, allele. (B) Locomotor activity measured for control adult flies (*THD1/+* and *FMRFaR^RNAi^/+*) were not different from that observed with *THD1>FMRFaR^RNAi^* flies (n=25, p>0.05, Mann-Whitney U-test).

**S2 Fig.**
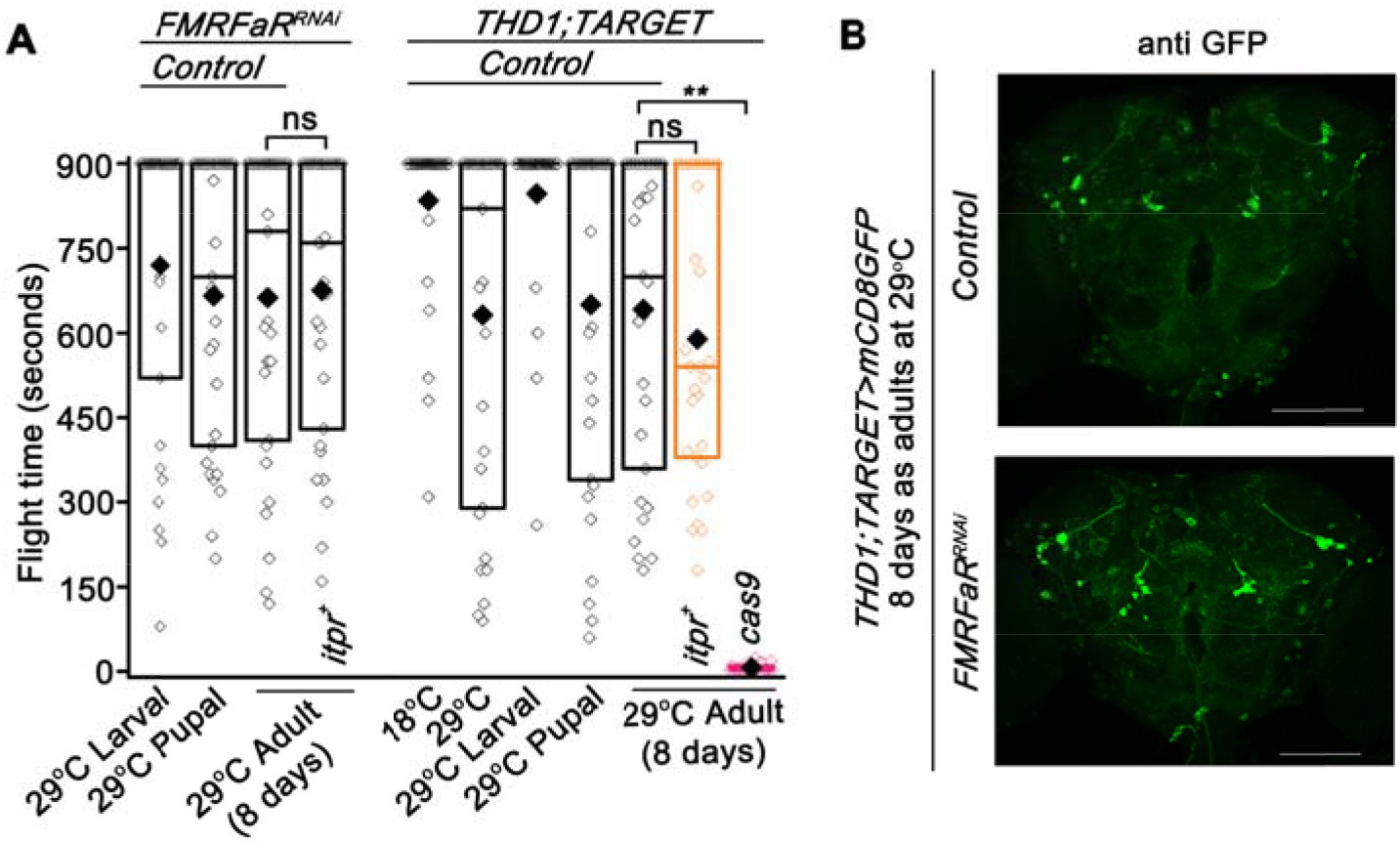
Behavior characterization under control conditions. (A) Genotypic controls for the TARGET experiment (*FMRFaR^RNAi^/+* and *THD1;TubGAL80^ts^ Control*) tested for flight under various temperature shift conditions. Other controls in the graph include *itpr^+^/+;FMRFaR^RNAi^/+* (4^th^ bar from the left in black), *THD1;TubGAL80^ts^>itpr^+^* (orange bar) and *THD1;TubGAL80^ts^>cas9* (pink bar). Comparisons are shown by horizontal lines (n=30, **p<0.01, ns – not significant; Mann-Whitney U-test). (B) Immunohistochemical staining of 8 day old adult brains using anti-GFP antibody in control (above; *THD1;TubGAL80^ts^>mCD8GFP* Control) and *FMRFaR* knockdown (below; *THD1;TubGAL80^ts^>mCD8GFP;FMRFaR^RNAi^*) conditions. Scale bars represent 100 μm.

**S3 Fig.**
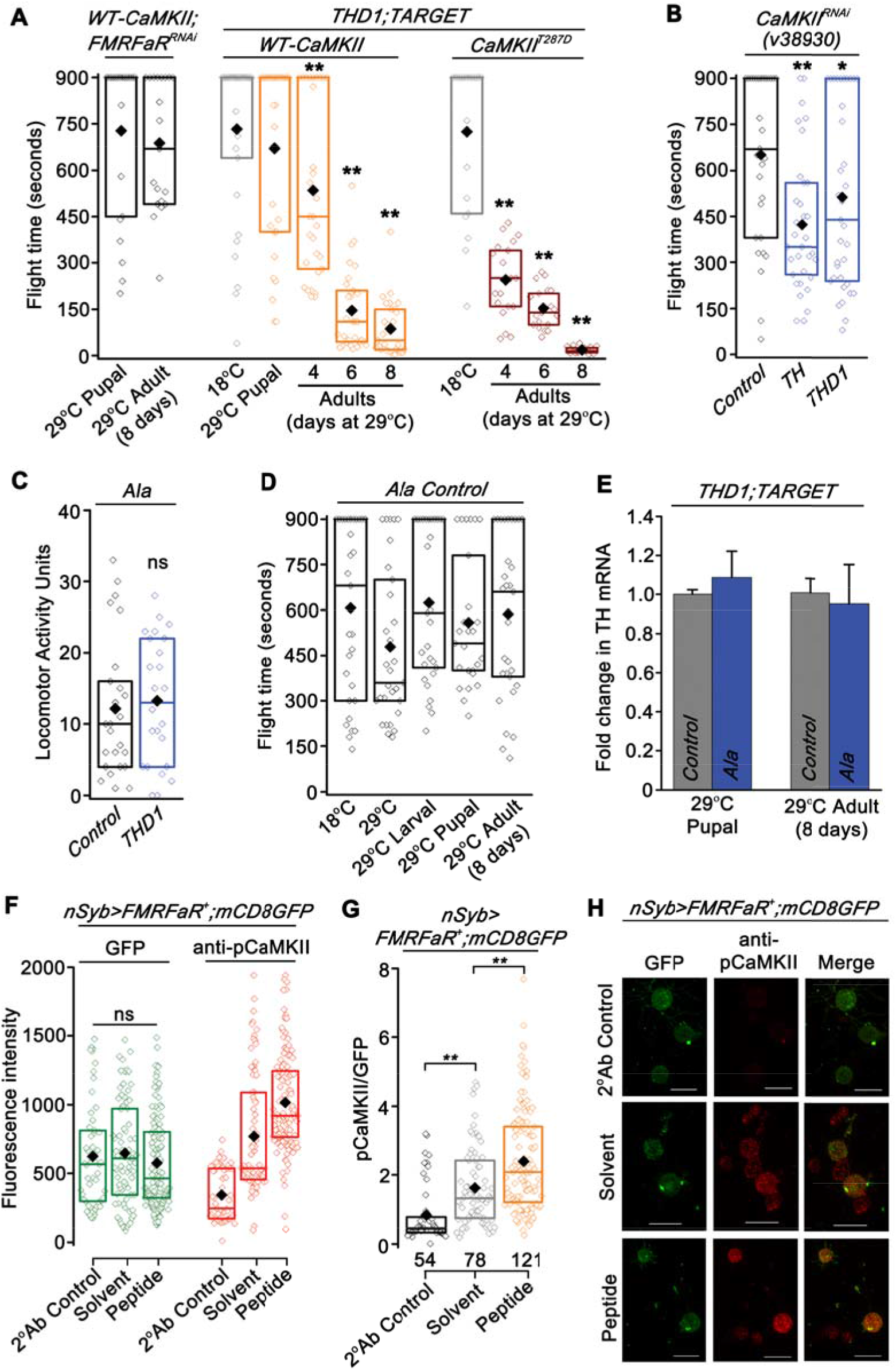
Behavioral and molecular characterization of CaMKII and *Ala* strains under various behavioral paradigms. (A) Flight durations observed for a strain used in Fig 4A (*WT-CaMKII/+;FMRFaR^RNAi^/+*). Flight bout durations measured for flies with expression of *WT-CaMKII (THD1;TubGAL80^ts^>WT-CaMKII*) or a constitutively active form of CaMKII, *CaMKII^T287D^ (THD1;TubGAL80^ts^>CaMKII^T287D^*), in *THD1* marked neurons under various temperature shift conditions as compared to the 18°C condition (nɥ20, **p<0.01, Mann-Whitney U-test). (B) Flight bout durations observed with knockdown of *CaMKII (CaMKII^RNAi^*) in dopaminergic neurons (n≥30, *p<0.05, **p<0.01, Mann-Whitney U-test). (C) Locomotor activity of control adult flies and flies with *THD1* driven *Ala* expression (*Ala/+* and *THD1>Ala;* n=25, p>0.05, ns - not significant; Mann-Whitney U-test). (D) Flight bout durations observed with an *Ala* control strain under the various temperature shift conditions (*Ala/+;* n≥20). (E) Quantitative PCR showing normalized mean fold change (± SEM) in *TH* transcripts under conditions of *Ala* expression in *THD1* neurons, either in pupae or adults (*THD1;TubGAL80^ts^ Control* and *THD1;TubGAL80^ts^>Ala;* n≥3, p>0.05, unpaired t-test). (F) Box plot showing background subtracted fluorescence intensity values of GFP and anti-pCaMKII under various experimental conditions. Cells with GFP intensities ranging from 0-1500 units were chosen for analysis and their distributions were not different in the three conditions (*nsyb>mCD8GFP;FMRFaR^+^;* p>0.05, ns - not significant; Mann-Whitney U-test). (G) Box plot showing the ratio of pCaMKII/GFP fluorescence intensity in the three experimental conditions. Peptide stimulation resulted in significantly higher pCaMKII/GFP ratios as compared to addition of the solvent (**p<0.01, Mann-Whitney U-test). (H) Representative images of cells showing endogenous GFP and staining for pCaMKII in the three treatment groups. Scale bar represents 10 μm.

**S4 Fig.**
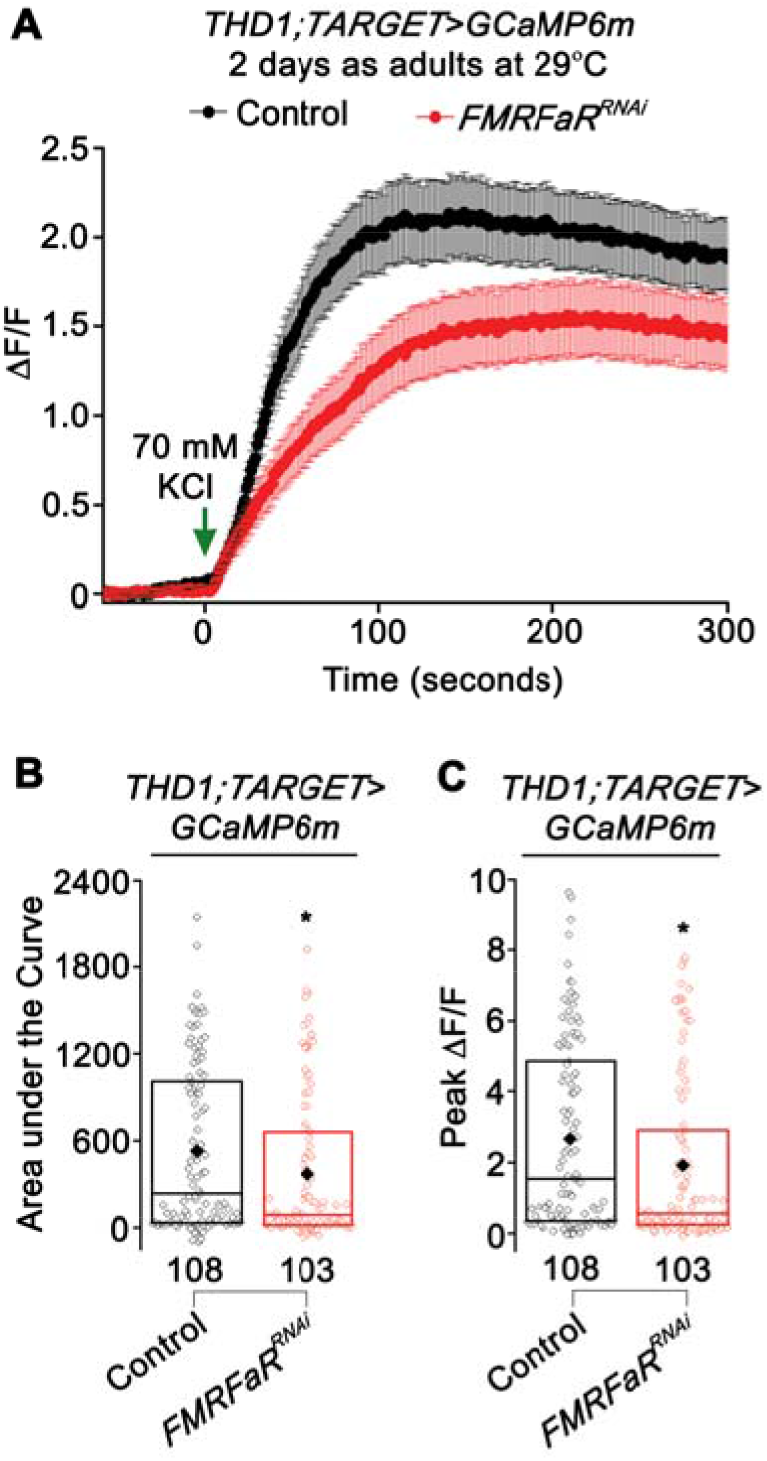
GCaMP6m imaging of *THD1* neurons after *FMRFaR* knockdown for 2 days in adults. (A) GCaMP6m traces showing normalized mean response (±SEM) of *THD1* neurons to KCl in 2 day old adult brains of the indicated genotypes, *THD1;TubGAL80^ts^>GCaMP6m, Control* in black; *THD1;TubGAL80^ts^>GCaMP6m;FMRFaR^RNAi^*, in red. (B) Area under the curve and (C) Peak ΔF/F quantified from (A). Numbers below each box plot indicate total number of cells imaged (*p<0.05; Mann-Whitney U-test).

**S5 Fig.**
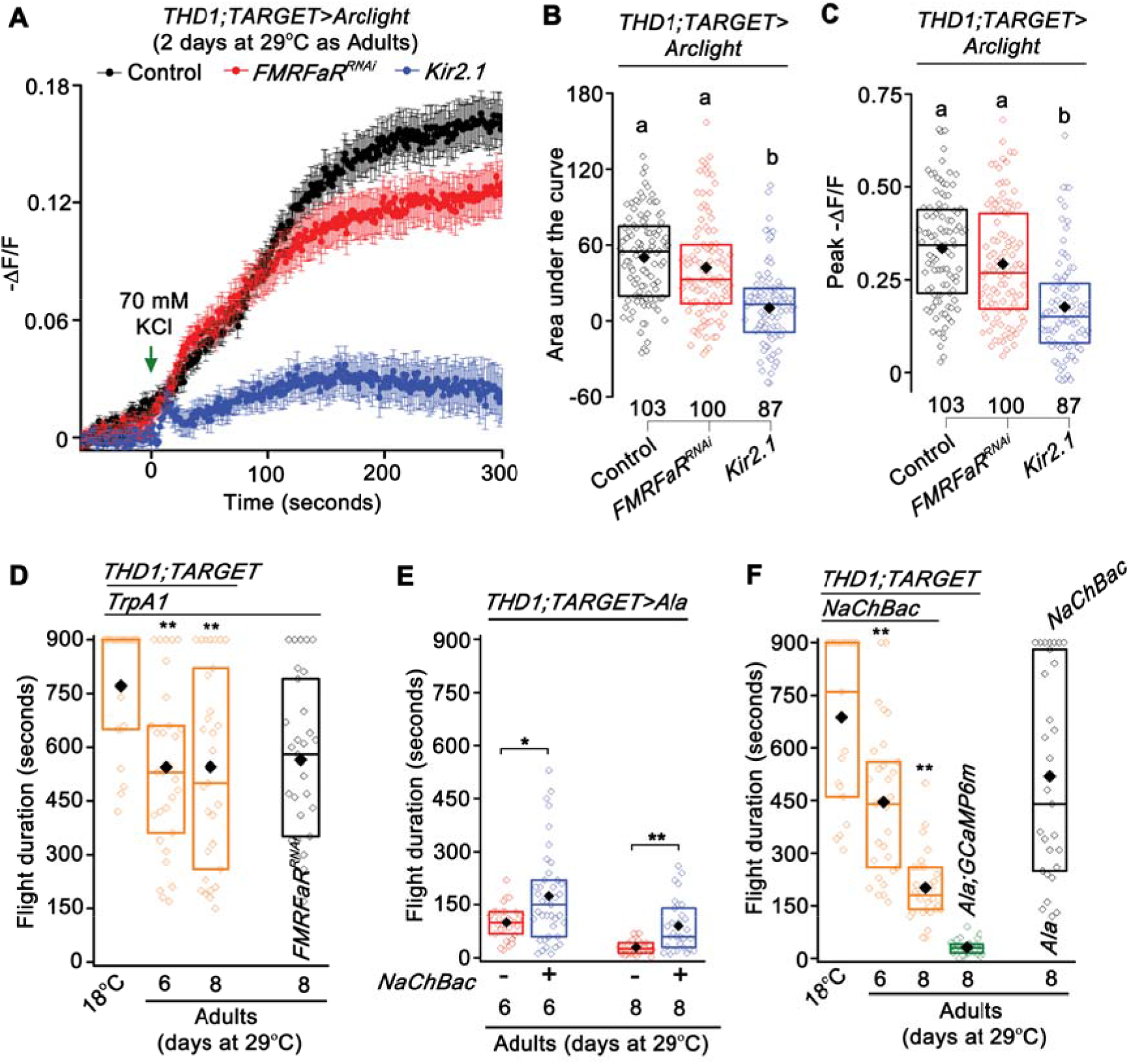
Cellular and behavioral characterization under control conditions. (A) Mean – ΔF/F traces observed in 2 day old adult dopaminergic neurons in response to KCl stimulation, where *THD1* neurons express the *Arclight* transgene in the indicated genotypes, *THD1;TubGAL80^ts^> Arclight, Control* in black; *THD1;TubGAL80^ts^>Arclight;FMRFaR^RNAi^*, in red, *THD1;TubGAL80^ts^>Arclight;Kir2.1* in blue. (B) Area under the curve and (C) Peak – (ΔF/F) calculated from (A). Expression of the hyperpolarizing channel, *Kir2.1*, significantly reduced the response to a depolarizing KCl stimulus. Numbers below each box plot indicate total number of cells imaged (One-way ANOVA followed by post-hoc Tukey’s test; the same alphabet above each bar represents statistically indistinguishable groups; different alphabet represents p<0.01). (D) Flight durations observed for flies expressing just the *TrpA1* transgene in adult *THD1* neurons as compared to 18°C control (*THD1;TubGAL80^ts^>TrpA1;* n≥20, **p<0.01, Mann-Whitney U-test). Flight duration of a control genotype used in Fig 6E (*TrpA1/+;FMRFaR^RNAi^/+* - black bar). (E) Rescue of flight deficits observed upon expression of *NaChBac*, in the background of CaMKII inhibition (Ala) in *THD1* neurons in 6 and 8 day old adults (*THD1;TubGAL80^ts^>Ala;NaChBac* compared to *THD1;TubGAL80^ts^>Ala;* n≥30, *p<0.05, **p<0.01, Mann-Whitney U-test). The rescues are presumably due to increased membrane excitability. (F) Flight bout durations observed with expression of just the *NaChBac* transgenes in *THD1* neurons under control condition (18°C) and for 6 or 8 days as adults (*THD1;TubGAL80^ts^>NaChBac;* orange bars; n≥20, **p<0.01, Mann-Whitney U-test). Flight times observed with expression of a *GCaMP6m* transgene in the background of *Ala* expression in adult *THD1* neurons (*THD1;TubGAL80^ts^>Ala;GCaMP6m;* green bar compared to *THD1;TubGAL80^ts^>Ala*, 8 days as adults at 29°C shown in Fig 4C are not significantly different; n≥20, p>0.05, Mann-Whitney U-test). Flight duration of a control genotype used in Fig 6E (*Ala/+;NaChBac/+* - black bar).

**S6 Fig.**
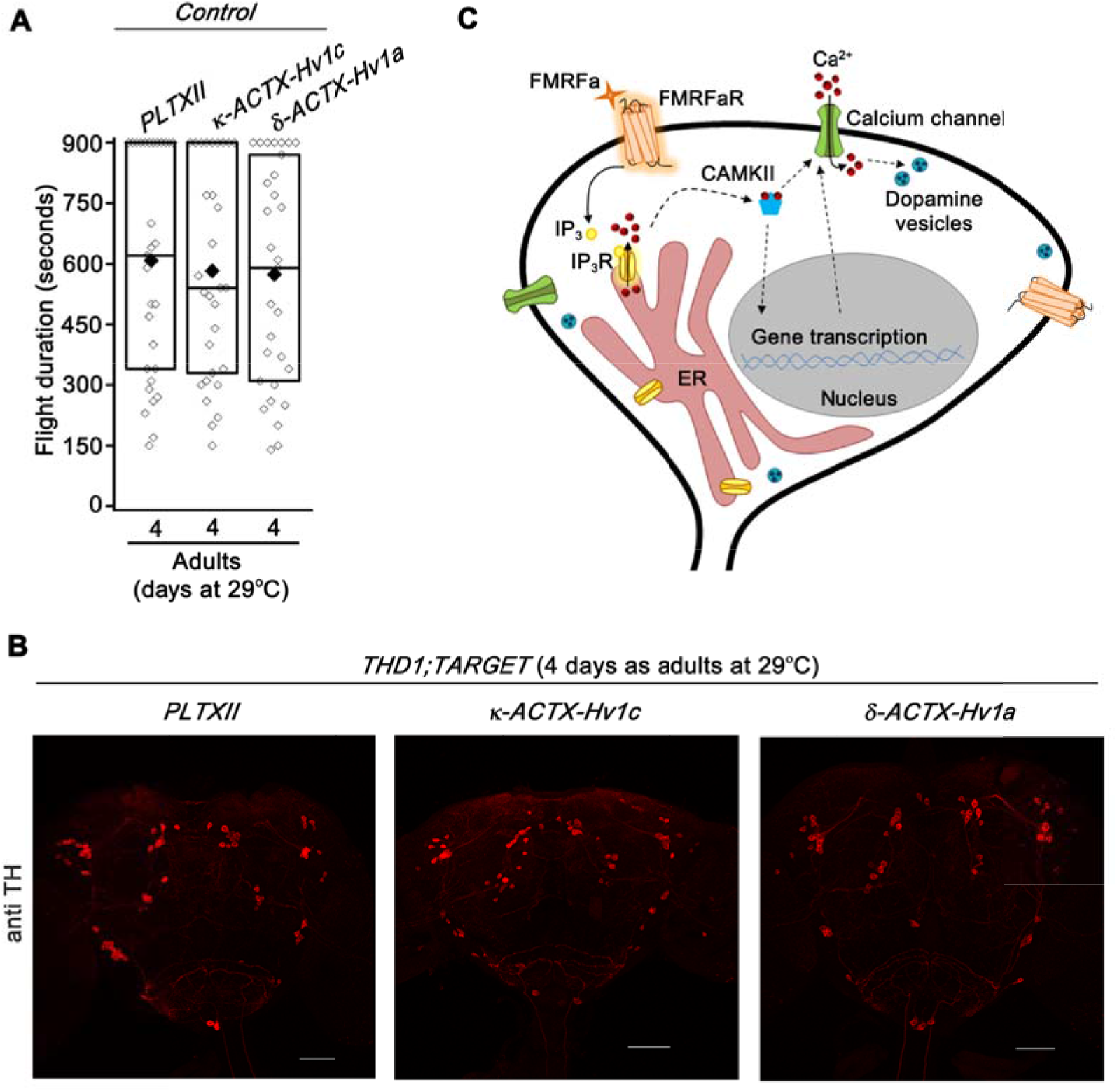
Behavioral and cellular characterization with channel toxin transgenes. (A) Flight times observed for controls of the three channel toxin transgenes (*PLTXII/+, κ-ACTX-Hv1c/+ and δ-ACTX-Hv1a/+;* n≥30, Mann-Whitney U-test). (B) Anti-TH immunostaining of 4 day old adult brains expressing the various channel toxin transgenes (*THD1;TubGAL80^ts^>PLTXII, THD1; TubGAL80^ts^>κ-ACTX-Hv1c and THD1;TubGAL80^ts^>δ-ACTX-Hv1a*). Neurons in various dopaminergic clusters were intact. Scale bars represent 100 μm. (C) Proposed signaling mechanism in dopaminergic neurons for regulation of flight bout durations by FMRFaR. The model illustrates that FMRFaR stimulation and downstream IP_3_R-mediated Ca^2+^ release could possibly activate CaMKII. Our data suggest that the FMRFaR and CaMKII are required in dopaminergic neurons for optimal membrane excitability. The effect on membrane excitability maybe by direct modification of membrane channels or by regulation of their expression levels. We further predict that this excitability is required for exocytosis of dopamine containing synaptic vesicles. Schematics are not drawn to scale.

